# Modification of regulatory tyrosines biases human Hsp90α for interaction with cochaperones and clients

**DOI:** 10.1101/2024.06.25.600625

**Authors:** Yuantao Huo, Rishabh Karnawat, Lixia Liu, Robert A. Knieß, Maike Gross, Xuemei Chen, Matthias P. Mayer

## Abstract

The highly conserved Hsp90 chaperones control stability and activity of many essential signaling and regulatory proteins including many protein kinases, E3 ligases and transcription factors. Thereby, Hsp90s couple cellular homeostasis of the proteome to cell fate decisions. High-throughput mass spectrometry revealed 178 and 169 posttranslational modifications (PTMs) for human cytosolic Hsp90α and Hsp90β, but for only a few of the modifications the physiological consequences are investigated in some detail. In this study, we explored the suitability of the yeast model system for the identification of key regulatory residues in human Hsp90α. Replacement of three tyrosine residues known to be phosphorylated by phosphomimetic glutamate and by non-phosphorylatable phenylalanine individually and in combination influenced yeast growth and the maturation of 7 different Hsp90 clients in distinct ways. Furthermore, wild-type and mutant Hsp90 differed in their ability to stabilize known clients when expressed in HepG2 *HSP90AA1*^−/−^ cells. The purified mutant proteins differed in their interaction with the cochaperones Aha1, Cdc37, Hop and p23 and in their support of the maturation of glucocorticoid receptor ligand binding domain *in vitro*. *In vivo* and *in vitro* data correspond well to each other confirming that the yeast system is suitable for the identification of key regulatory sites in human Hsp90s. Our findings indicate that even closely related clients are affected differently by the amino acid replacements in the investigated positions, suggesting that PTMs could bias Hsp90’s client specificity.

## Introduction

The 90 kDa heat shock proteins (Hsp90s) are highly conserved and essential molecular chaperones in all eukaryotic cells [1]. Hsp90s are especially known for chaperoning some 300 substrate proteins, called clients, in a late folding, near-native state. Among these clients are about 60% of all human protein kinases, about 30% of the E3 ligases, and many transcription factors, including steroid hormone receptors [2]. As many oncogenes like Raf, Src and receptor tyrosine kinases, and tumor suppressors like p53, BRCA1 and BRCA2 are among the Hsp90 clients, Hsp90 is considered to be a prime target for cancer therapy [3, 4].

Hsp90s consist of an N-terminal nucleotide binding domain that is connected via an intrinsically disordered charged linker of different lengths to a middle domain, followed by a dimerization domain and a second intrinsically disordered region of different lengths. All eukaryotic cytosolic Hsp90s end with the highly conserved EEVD motif. Structural studies revealed that Hsp90 can assume several different conformations from a wide-open V-shaped to a closed intertwined conformation in which the N-terminal domains of the two Hsp90 protomers are docked onto each other and onto the middle domain [5–8]. The Hsp90 chaperone cycle was first elucidated for steroid hormone receptors [9]. Inactive steroid hormone receptors are first bound by Hsp70 with the help of its J-domain protein (JDP) cochaperone and are subsequently transferred to Hsp90. This transfer is assisted by the cochaperone Hop/Sti1, which binds like a scaffold to the C-terminal EEVD motifs of cytosolic Hsp70s and Hsp90s. The cochaperone p23 displaces Hop and Hsp70 and assist the transition to the mature Hsp90 client complex [10]. Kinase clients are first bound by the specific cochaperone Cdc37 and subsequently transferred to Hsp90 [11, 12]. Aha1 seems to play a role in the chaperoning of kinases by Hsp90 [13, 14]

As Hsp90 interacts with late folding intermediates and near-native proteins the question arises how Hsp90 can recognize so many different clients that are unrelated in sequence and structure. One answer might be the extended surface of Hsp90s that contains hydrophobic as well as hydrophilic and charged patches and offers many low affinity interaction sites to which different clients can bind in distinct ways [15–20]. The large conformational flexibility might contribute to client binding, allowing Hsp90 to adapt to many protein folds. In addition, Hsp90 interacts with more than 30 cochaperones that target the clients to Hsp90 and might stabilize the Hsp90-client complex [21–23].

Hsp90 is modified by a large number of post-translational modifications (PTMs). There are

two isoforms of Hsp90 in the cytosol of human cells, Hsp90α and Hsp90β, encoded by the genes HSP90AA1 and HSP90AB1, and the PhosphoSitePlus® database (28.05.2024) lists 74 and 66 phosphorylation sites and 41 and 34 acetylation sites for Hsp90α and Hsp90β, respectively (**Fig. S1A**). These sites were detected by high throughput mass spectrometry experiments and only few were investigated to some degree (e.g. [11, 24–30]). Therefore, the physiological significance of most of the PTMs is currently unexplored. Some of the modifications might be fortuitous without further consequences for the activity of Hsp90, some might be important for progression through Hsp90’s conformational cycle [11], some for client release as shown for arylhydrocarbon receptor [31] or protein kinase Cγ [32], and some might be important for client protein selection. It is therefore important to analyze all PTMs and also different combinations of them, since it is not known how different PTMs affect each other’s functions (e.g. priming phosphorylation). To do this site by site seems hardly feasible and a high-throughput method would be necessary. To explore this avenue, we selected three tyrosine sites in human Hsp90α, two of which had been analyzed previously to some degree [11, 26]. We used the genetically tractable organism yeast as *in-vivo*-model system to investigate the effect of phosphomimetic and non-phosphorylatable amino acid replacement on growth of yeast and on seven Hsp90 clients, five steroid hormone receptors and two kinases. In HepG2 *HSP90AA1*^−/−^ cells, we tested the ability of wild-type and mutant Hsp90 to stabilize five known Hsp90α client proteins. We complemented the *in-vivo*-findings with *in-vitro*-experiments using purified proteins. We were particularly interested in the question whether phenylalanine and glutamate are suitable surrogates for tyrosine and phosphorylated tyrosine, respectively, and whether alterations at multiple sites act synergistic, additive, or antagonistic. Our data indicate that all three investigated sites are regulatory hotspots that influence even closely related clients in distinct manners.

## Material and Methods

### Yeast experiments

The *S. cerevisiae* strain MPMH2458 (MAT**a**, *ade2-101oc, his2Δ200, leu2-3, lys2-801^am^, trp1-289, ura3-52, Δpdr5::Blank, Δhsp82::KanMX, Δhsc82::KanMX with plasmid* YEplac195(2µ-HSC82-URA3) used in this study is a derivation of DP533 (courtesy of D. Picard [33]). Yeast genetics and assays were performed according to standard procedures [34]. For details see supplemental information. For expression of hHsp90α or its mutants in *S. cerevisiae* an expression cassette was integrated into the LYS2 locus by homologous recombination. The expression cassette contained the *Klyvermyces lactis* LEU2 marker flanked on both sites as inverted repeats by the HSP90AA1 gene fused N-terminally to the FLAG-tag sequence under the control of the strong TDH3 promoter (**Fig. S1B**). For the reporter assay, the yeast strain MPMH2458 with integrated wild-type or mutant HSP90AA1 expression cassettes were co-transformed with plasmids p413GPD-GR, p413GPD-MR, p413GPD-AR, or p413GPD-PR, encoding the hormone receptors, with plasmid pUCΔSS-26X, encoding the steroid response element responsive to GR, MR, AR and PR, or with p413GPD-ER, encoding estrogen receptor, together with pΔsERE, encoding the estrogen response element (courtesy of J. Buchner [35]). For the v-Src maturation assay, pMPM0484, the empty vector was used as a negative control. For a detailed plasmid list see supplemental **Table S1**.

### Protein purification

All proteins, except Glucocorticoid receptor ligand binding domain (GRLBD), were recombinantly expressed as N-terminal His6-SUMO fusions [64] and purified by Ni^2+^-affinity chromatography, anion exchange chromatography and size exclusion chromatography. The tetra-cysteine variants of the cochaperones had the sequence “ACCPGCCSGG” at the N-terminus (following the SUMO-tag) or at the C-terminus of the protein. Human GRLBD (GR(521-777)) with the stabilizing amino acid replacement F602S was expressed as N-terminal fusion to maltose binding protein [43]. For details see supplemental information.

### Binding Affinity Measurements

All cochaperones had at their N- or C-terminus a tetra-cysteine tag (ACCPGCCSGG) and were labeled with FlAsH-EDT2™. For the fluorescence anisotropy experiments, the labeled cochaperone (∼100 nM) was added to a serial dilution of Hsp90 and the samples pipetted into a black Corning® Low Volume 384-well plate. In the case of p23, Hsp90 was first incubated with 4 mM AMPPNP at room temperature for 20 min before the labeled cochaperone was added. The samples were measured on a CLARIOstar plate reader (BMG Labtech). The anisotropy values were fitted to the quadratic solution of the law of mass action to obtain the KD. For more details see supplemental information.

### Differential Scanning Fluorimetry

The Hsp90 variants at a concentration of 3 µM in 25 mM HEPES/KOH pH 7.5, 150 mM KCl, 5 mM MgCl2, 10% glycerol, and 2 mM DTT were heated in a LightCycler® 480 Multiwell white 384-well plate in a Roche LightCycler® 480 II from 20 to 80°C in the presence of SYPRO™ Orange (Merck) diluted 1:160.

### In vitro GR refolding assay

Hsp90, Hsc70, p23, Hop, Ydj1, and MBP-GRLBD were thawed and centrifuged for 30 min at 20,000 g at 4°C. The buffer was exchanged for refolding buffer (40 mM HEPES/KOH pH 7.5, 50 mM KCl, 5 mM MgCl2, and 2 mM DTT) using Zeba™ Spin Desalting Columns. GRLBD was desalted twice using a Zeba™ Spin Desalting Column to remove the unbound dexamethasone. A mixture containing 1 µM GRLBD, 12 µM Hsc70, 2 µM Ydj1, and 5 mM ATP was incubated at 42°C for 10 min to further unfold GRLBD. Subsequently, 12 µM Hop, 12 µM p23, and 12 µM Hsp90 and 100 nM of fluorescein-labeled dexamethasone (F-dex; Thermo Fisher Scientific) were added, the samples were transferred into a Corning® Low Volume 384-well black plate and measured at 25°C in a CLARIOstar Plate Reader (BMG Labtech). A heated sample of GRLBD alone served as negative control and GRLBD without heat treatment served as the natively folded control. Radicicol was used as a control to test for the influence of Hsp90 in the maturation process. A sample containing only F-dex and buffer was used as blank and subtracted from all fluorescence anisotropy values.

### Cell culture and reagents

HepG2 (SCSP-510) cells were purchased from the Cell Bank, Chinese Academy of Medical Sciences (Shanghai, China). All cells were thawed upon arrival, expanded, and stored in a liquid nitrogen tank. The cells were confirmed to be free of mycoplasma contamination with MycAwayTM-Color One-Step Mycoplasma Detection Kit (40612ES60, Yeasen, Shanghai, China). HepG2, were cultured in Dulbecco’s modified Eagle’s medium (DMEM, Gibco, USA) with 10 % fetal bovine serum (Gibco, USA) at 37°C, 5% CO2.

### Hsp90α-knockout construction

CRISPR-Cas9 plasmids for Hsp90α knockout were purchased from iGeneBio (Guangzhou, China); Hsp90α sgRNA sequences were sgRNA-F:5’-TTCTTGGGTAGTTTGCAG-3’, sgRNA-R: 5’-CGCCATTCGCCATTCAGG-3’. HepG2 Cells were transfected with the sgRNA plasmid for Hsp90α with Lipofectamine 3000 (Invitrogen, CA, USA). After 48 h, cells were collected, and incubated in selection medium containing 800 μg/mL G418 for 2–3 weeks. The knockout clones were confirmed by immunoblotting.

### Total protein lysates and Western Blot

Cells were lysed in RIPA lysis buffer (P0013B, Beyotime Biotechnology, China) (50 mM Tris (pH 7.4), 150 mM NaCl, 1% Triton X-100, 1% sodium deoxycholate, 0.1% SDS) containing protease inhibitor (Pierce, MA, USA). Western blot analysis was performed as described previously [36, 37]. For primary antibodies see supplemental **Table S2**. The secondary antibodies (Donkey anti-Mouse/Rabbit IRDye 680/800, Abcam, USA,1:20000) were from Abcam. Western blots were scanned using Li-COR Odyssey, and band intensities were analyzed by ImageJ.

## Results

### Human Hsp90α variants differ in their ability to complement Hsp90 deficiency in yeast

To establish a methodology to thoroughly characterize the potential impact of tyrosine phosphorylation at specific sites in human Hsp90 and to explore possible cross-talk between different phosphorylation events, we chose three sites in the N-terminal domain of human Hsp90α, Y38, Y61 and Y197, all of which had been found in multiple high throughput phosphoproteomics studies (see https://www.phosphosite.org/) (**Fig. 1A & B, Fig. S1A**). Two

**Figure 1:**
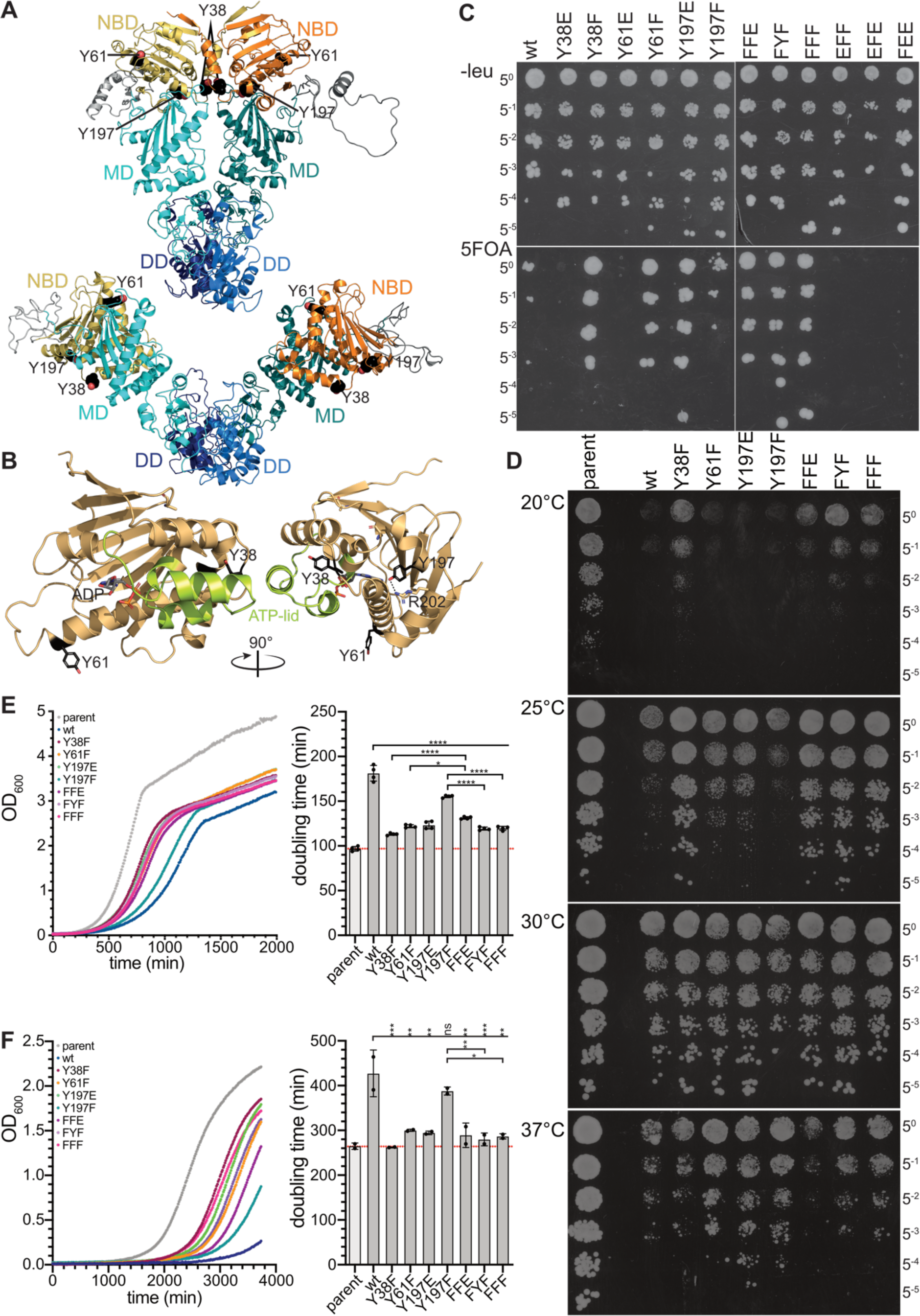
Phosphomimetic and non-phosphorylatable variants of human Hsp90α differentially support growth of *S. cerevisiae* cells lacking endogenous Hsp90. **A:** Cartoon representation of a homology model of the human Hsp90α dimer on the crystal structure of the closed conformation of yeast Hsp82 in complex with AMPPNP and sba1 (2CG9; [5]; upper panel) and on the crystal structure of the open conformation of apo *E. coli* HtpG (2IOQ; [61]; lower panel). Nucleotide binding domain (NBD), yellow and orange; middle domain (MD), cyan and teal; C-terminal dimerization domain (DD), blue and dark blue; tyrosine residues, Y38, Y61 and Y197, investigated in this study are shown as spheres in atom colors with carbon in black. **B:** Cartoon representation of the NBD of human Hsp90α in complex with ADP (1BYQ; [62]). Right panel rotated by 90° as indicated. So-called ATP-lid shown in green. Tyrosine residues, Y38, Y61 and Y197, investigated in this study are shown as sticks in atom colors with carbon in black. Potential polar interactions shown as dashed lines. **C:** Complementation of lethality phenotype of the lack of endogenous Hsp90 in *S. cerevisiae* by wild-type hHsp90α or phosphomimetic and non-phosphorylatable variants. Plasmid shuffling assay. The expression cassette shown in Fig. S1B encoding two wild-type or mutant genes of hHsp90α (*HSP90AA1*) was integrated into the *LYS2* locus of the *S. cerevisiae* strain MPMH2458 in which both Hsp90 encoding genes *HSP82* and *HSC82* had been deleted and which carried a 2µ-episomal plasmid expressing *HSC82* and *URA3*. Fivefold dilutions of an overnight culture containing the indicated hHsp90α variant were spotted onto selective media minus leucine (SD-leu; upper panel) or 5-fluoroorotic acid (5FOA; lower panel), selecting for the loss of the URA3 containing plasmid with the *HSC82* gene. FFE: Y38F,Y61F,Y197E; FYF: Y38F,Y197F; FFF: Y38F,Y61F,Y197F; EFF: Y38E,Y61F,Y197F; EFE: Y38E,Y61F,Y197E; FEE: Y38F,Y61E,Y197E; Loss of yHsc82 and hHsp90α levels of complementing strains were verified by immune blotting (see Fig. S1C). Shown is one of three independent experiments with similar results. **D:** Temperature sensitivity of yeast cells expressing wild-type or mutant human *HSP90AA1* as sole source for Hsp90. Fivefold dilutions of overnight yeast cultures expressing high levels of yHsc82 (parental) or wild-type or mutant hHsp90α were spotted on YPD plates and grown at the indicated temperatures for 72 h (growth after 48 h is shown in Fig. S1D). **E & F:** Growth curves for yeast stains containing yHsc82 or wild-type and mutant hHsp90α in YPD (E, left panel) or SCGE (selective complete with 3% glycerol, 2% ethanol; F, left panel) (for more clarity see Fig. S1E & F). Right panels, doubling times calculated from the growth curves by fitting an exponential function to the OD values below 0.5. Shown are mean and standard deviation. Statistical significance was tested by ordinary one-way ANOVA with Šidák’s multiple comparison; ns, not significant; *, p < 0.05; **, p < 0.01; ***, p < 0.001; ****, p < 0.0001. All hHsp90α containing strains were compared to the strain containing wild-type hHsp90α and individual pairs were compared as indicated. The human Hsp90α variants Y38E and Y61E do not support growth of the Δhsc82Δhsp82 yeast strain, indicating that a negative charge in position 38 and 61 of hHsp90α does not allow the maturation of essential clients in yeast. In contrast, the hHsp90α variants Y38F, Y61F and Y197E complemented the lack of Hsp90 in yeast even better than wild-type hHsp90α and Y197F as demonstrated in higher plating efficiency and larger colonies on plates (Fig. 1C) and in higher growth rates in liquid media (Fig. 1E, Fig. S1E). These differences in growth were observed in media with the fermentable carbon source glucose as well as in media with the non-fermentable carbon source glycerol (3%) plus ethanol (2%) (Fig. 1F, Fig. S1F).

of the sites, Y38 and Y197, had been investigated previously to some extend [11, 26, 38], whereas Y61 phosphorylation has not been investigated in greater details to our knowledge. To test the chaperone activity of the phosphomimetic (Y-to-E) and non-phosphorylatable (Y-to-F) variants of human Hsp90α in a complex cellular system without endogenous Hsp90, we resorted to the *Saccharomyces cerevisiae* model system. It had been shown previously that human Hsp90 can complement the lethal phenotype of deletion of the Hsp90-encoding genes in budding yeast [33] and that it is suitable for studying the maturation of yeast and human clients [35, 39, 40]. We utilized a yeast strain that had both endogenous Hsp90-encoding genes, *HSP82* and *HSC82*, deleted but expressed *HSC82* extra-chromosomally on a URA3-containing rescue plasmid. This strain was transformed with integration constructs expressing human *HSP90AA1* wild-type and mutant variants (**Fig. S1B**). After transformation/integration, we selected for loss of the URA3-plasmid by plating on 5-fluoroorotic acid (5FOA) (“plasmid/gene shuffling”), and thereby for the loss of the *yHSC82* gene, leaving strains that contain hHsp90α wild-type or mutant proteins as sole source of Hsp90 (**Fig. 1C**). This replacement was confirmed by immunoblotting (**Fig. S1C**).

The Hsp90 variants Y38E and Y61E did not complement yeast growth. In contrast, the strains containing the non-phosphorylatable variants Y38F and Y61F or the phosphomimetic Y197E grew significantly better than the strain containing wild-type Hsp90. For the surprising phenotype of Y38F and Y61F there are two possible explanations: (1) phenylalanine has a gain of function over tyrosine (e.g. due to its greater hydrophobicity); (2) Although wild-type and mutant proteins are expressed at similar levels in yeast cells, a subpopulation of wild-type Hsp90 is inactive due to phosphorylation in yeast by the dual-specificity serine/threonine-tyrosine kinase Swe1, the only tyrosine kinase in yeast. It had been shown previously that wild-type yeast and human Hsp90 can be phosphorylated at a single tyrosine residue (Y38) in yeast cells [26]. To test this hypothesis, we isolated Hsp90α from yeast cells after plasmid shuffling using the N-terminal FLAG-tag and performed phosphopeptide enrichment coupled with mass spectrometry. Although we detected several serine and threonine phosphorylated proteolytic peptides of hHsp90α, we did not detect any tyrosine phosphorylated peptides, suggesting that wild-type hHsp90α is not phosphorylated in any of the three tyrosine residues investigated in this study (Y38, Y61, or Y197) to any detectable extent. Therefore, the reason for the slow growth phenotype of the strain containing wild-type hHsp90α, as compared to either Y38F or Y61F containing strain, may not be due to Y38 or Y61 phosphorylation.

To our knowledge, no study has been conducted so far that examined whether the combination of multiple phosphomimetic or non-phosphorylatable amino acid replacements in human Hsp90α could have synergistic or neutralizing effects. To test any synergistic effects, we combined variants that are more growth-supportive to generate the variants FFE (Y38F,Y61F,Y197E), FYF (Y38F,Y197F), and FFF (Y38F,Y61F,Y197F). To test antagonistic and compensatory effects, we combined the Y38E and Y61E replacements with two more growth-supportive replacements, generating EFF (Y38E,Y61F,Y197F), EFE (Y38E,Y61F,Y197E) and FEE (Y38F,Y61E,Y197E). Combining the more growth-supportive replacements with each other did not result in an additional benefit for yeast growth (**Fig. 1C**, **Fig.1E** right panel, **Fig. 1F, Fig. S1E & F**) and combining more growth-supportive amino acid replacements with an amino acid replacement that does not support growth, did not result in the rescue of the yeast strain (**Fig. 1C**).

Interestingly, different stains displayed different types or levels of temperature sensitivity (**Fig. 1D, Fig. S1D**). When incubated at low temperatures, cells containing an Hsp90α variant with the Y38F amino acid replacement grew much better than strains containing wild-type Hsp90α, Y61F, Y197E or Y197F variants. However, the latter strains tolerated higher temperatures better than the ones that have the Y38F replacement alone or in combination with other replacements.

### Hsp90 mutants differ in their ability to support maturation of diverse clients

Hsp90 is known to chaperone a wide range of clients, including kinases, E3 ligases, transcription factors, and receptors [2, 41]. To investigate the impact of phosphomimetic and non-phosphorylatable amino acid replacements in human Hsp90α on the maturation and activity of Hsp90 clients, we measured the transcriptional activity of the five highly homologous human steroid hormone receptors (SHRs), glucocorticoid receptor (GR), mineralocorticoid receptor (MR), androgen receptor (AR), progesterone receptor (PR), and estrogen receptor (ER) using a *lacZ-*based reporter assay. The activity of the yeast serine/threonine kinase Ste11, another known Hsp90 client [42], we determined using a *lacZ* reporter construct responsive to the transcription factor Ste12 that is activated through Ste11-Ste7-Fus3/Kss1 kinase cascade [42]. In addition, we assessed the activity of the oncogenic tyrosine kinase v-Src that inhibits yeast growth when overexpressed and activated by Hsp90 [40].

For the five human steroid hormone receptors (SHRs), we measured the basal activity in the absence of hormone (w/o) and the induced activity in the presence of the respective hormone 11-Deoxycorticosterone (DOC), Dihydrotestosterone (DHT), progesterone, aldosterone, or β-estrogen. Striking differences in the induced activity of the different SHRs were observed (**Fig. 2A-E**). GR exhibited the highest activity in the presence of wild-type Hsp90α and Y197F, whereas in the presence of all other variants GR was less active. MR and AR had the highest activity in the presence of Y197F, whereas wild-type Hsp90 and the other Hsp90 variants were markedly less efficient to support maturation of these two SHRs. On the other side, PR was best activated by Y38F and significantly less by Y197F and Y197E, whereas wild-type Hsp90α and the other variants failed to activate PR. ER was matured well by wild-type Hsp90, Y38F and Y197F but not by the other Hsp90 variants. In the combination variants the negative effects seemed to dominate the competence of Hsp90 to activate its client proteins.

**Figure 2:**
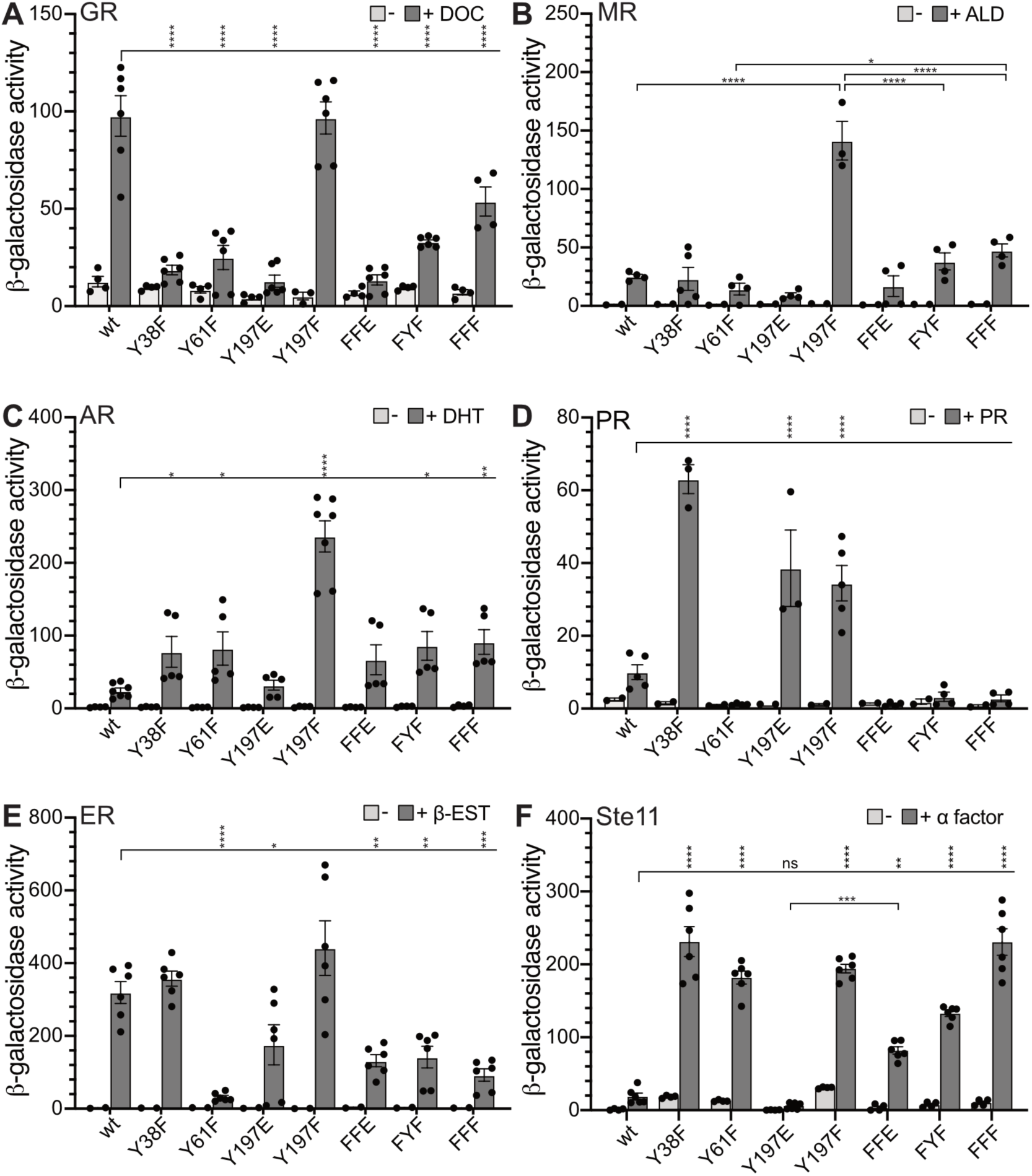
Phosphomimetic and non-phosphorylatable hHsp90 variants differ in their ability to assist maturation of Hsp90 clients in the yeast model system. **A-F**: *Δhsc82 Δhsp82* yeast cells containing wild type or mutant hHsp90 and expressing human glucocorticoid receptor (**A**), human mineralocorticoid receptor (**B**), human androgen receptor (**C**), human progesterone receptor (**D**), human estrogen receptor (**E**) and a corresponding response element driven *lacZ* reporter construct (**A-E**) or a Ste11/Ste12 responsive *lacZ* reporter construct (**F**) were grown at 30°C in the absence (light gray bars) of presence (dark gray bars) of 10 µM of the corresponding hormone or α-mating pheromone for 3 h and β-Galactosidase activity was determined as described in material and methods. DOC, deoxycorticosterone; ALD, aldosterone; DHT, dihydrotestosterone; PR, progesterone; β-EST, β-estradiol. FFE: Y38F,Y61F,Y197E; FYF: Y38F,Y197F; FFF: Y38F,Y61F,Y197F. Shown are mean and standard deviation. Statistical significance of differences relative to wild-type hHsp90 or between individual pairs of hHsp90 variants was tested by ordinary one-way ANOVA with Šidák’s multiple comparison; ns, not significant; *, p < 0.05; **, p < 0.01; ***, p < 0.001; ****, p < 0.0001.

Ste11 activity was well supported by Y38F, Y61F and Y197F but not by wild-type Hsp90α and Y197E (**Fig. 2F**). Combination of the three amino acid replacements that supported Ste11 maturation (FFF: Y38F,Y61F,Y197F) did not lead to increased activity, indicating that there is no synergistic or additive effect. Combination of Y197E with Y38F and Y61F resulted in an intermediate activity of Ste11, suggesting that the negative effect of the negative charge in position 197 could be partially compensated by the phenylalanine in positions 38 and 61.

To assess the impact of the Hsp90 variants on v-Src activity, we expressed *V-SRC* under the galactose-inducible GAL1 promoter on galactose containing plates or in galactose containing liquid media, and compared growth of the *V-SRC* expressing yeast strains with the growth of the isogenic strains harboring an empty vector. The parental strain containing *yHSC82* that shows no growth on a galactose plate and reduced growth in liquid media (**Fig. 3A-C**) was used as positive control. For the two Hsp90 variants Y197E and Y61F, no growth difference between the *V-SRC* expressing strain and the empty vector containing strain were observed on plate and in liquid culture, indicating the inability of these two variants to support maturation of v-Src. For wild-type human Hsp90α and the other variants differences between the *V-SRC* expressing strain and the empty vector strain were observed, albeit not as prominently as for the strain containing wild-type yHsc82. This indicates a reduced ability of human Hsp90α to support v-Src maturation as compared to yHsc82, which could be due to different expression levels or reduced ability to cooperate with the yeast cochaperones.

**Figure 3:**
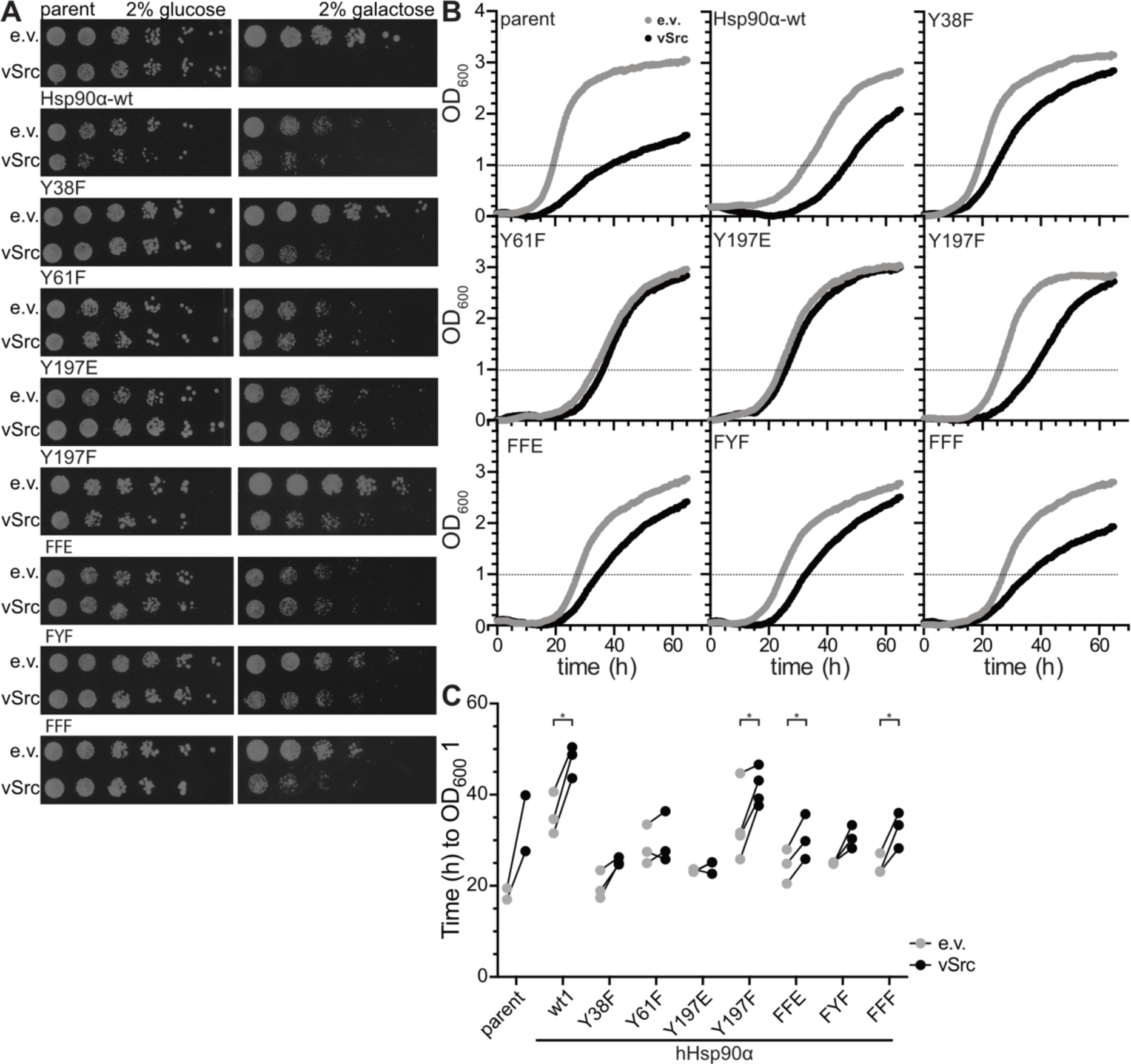
Phosphomimetic and non-phosphorylatable hHsp90 variants differ in their ability to assist maturation of v-Src in the yeast model system. **A**: overnight cultures of the *Δhsc82 Δhsp82* yeast strain expressing y*HSC82* (parent) or genes for wild-type or mutant hHsp90 (after 5FOA selection) and containing an empty vector (e.v.) or a vector expressing *V-SRC* under the control of a galactose-inducible promoter were spotted in 10-fold dilutions on YP-plates with 2% glucose or 2% galactose. FFE: Y38F,Y61F,Y197E; FYF: Y38F,Y197F; FFF: Y38F,Y61F,Y197F. One of two independent experiments with similar results is shown. **B**: Strains as in A were grown in liquid media containing 2% galactose and OD600 was measured (empty vector, e.v., light gray data points; *V-SRC*-expressing plasmid, black data points). One of two to four independent experiments is shown. **C**: Growth time. From the data shown in B and similar data of biological replicates the time was determined for each culture to reach an OD600 of 1. Statistical significance of differences was determined by paired t-tests. *, p < 0.05.

To assess the impact of the Hsp90 variants on clients in human cells, we transfected *HSP90AA1*^−/−^ HepG2 cells with plasmids expressing wild-type and mutant Hsp90α and determined the levels of the known Hsp90α clients DNA-dependent protein kinase catalytic subunit (DNA-PKcs), Nibrin (NBN), Bcl-2-associated transcription factor 1 (BCLAF1), RAC-alpha serine/threonine-protein kinase (AKT1), and Voltage-dependent anion-selective channel protein 1 (VDAC1) by immunoblotting (**Fig. 4**). DNA-PKcs and NBN levels were significantly higher in the presence of Y61E than in the presence of Y61F, suggesting that phosphorylation at Y61 stabilizes these Hsp90α clients. For other Hsp90α variants we did not observe significant differences for any of the clients.

**Figure 4:**
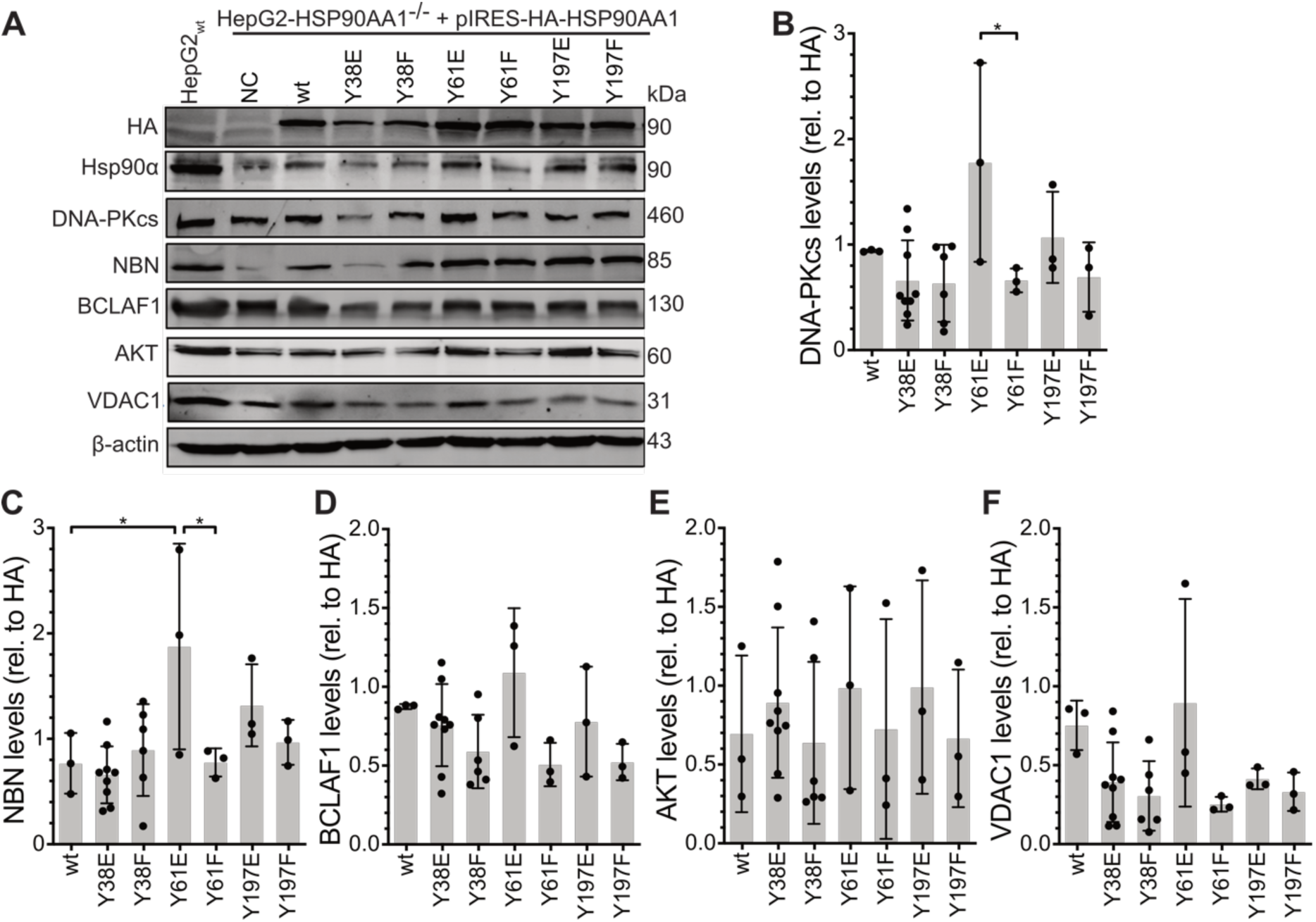
Ectopically expressed phosphomimetic and non-phosphorylatable hHsp90α variants in HepG2 *HSP90AA1*^−/−^ cells differ in their ability to chaperone different clients. **A**: Example immunoblots using HA, Hsp90α, DNA-PKcs, NBN, BCLAF1, AKT, VDAC1 and β-actin specific antisera. HepG2 *HSP90AA1*^−/−^ were transiently transfected with plasmids expressing the indicated HA-tagged Hsp90α variant. NC, vector only control. **B**-**F**: Quantification of immunoblots shown in A and similar blots of independent experiments. Shown are mean and standard deviation. Statistical significance was tested with ordinary one-way ANOVA and Šidák’s multiple comparison; *, p < 0.05.

*Amino acid replacements affect the conformation of human Hsp90α*

To elucidate the molecular mechanism of how the amino acid replacements affect the chaperone function of Hsp90α, we purified the recombinant glutamate and phenylalanine variants and analyzed them *in vitro*. We first assessed the thermodynamic stability of the proteins by differential scanning fluorimetry (DSF) at temperatures between 20 and 80°C using SYPRO^®^ Orange (**Fig. 5A**). For comparison and to interpret the observed transitions, we analyzed the individual domains (N-terminal nucleotide binding domain (N), the middle domain (M) and the C-terminal dimerization domain (C)) and, in addition, N-M and M-C constructs.

**Figure 5:**
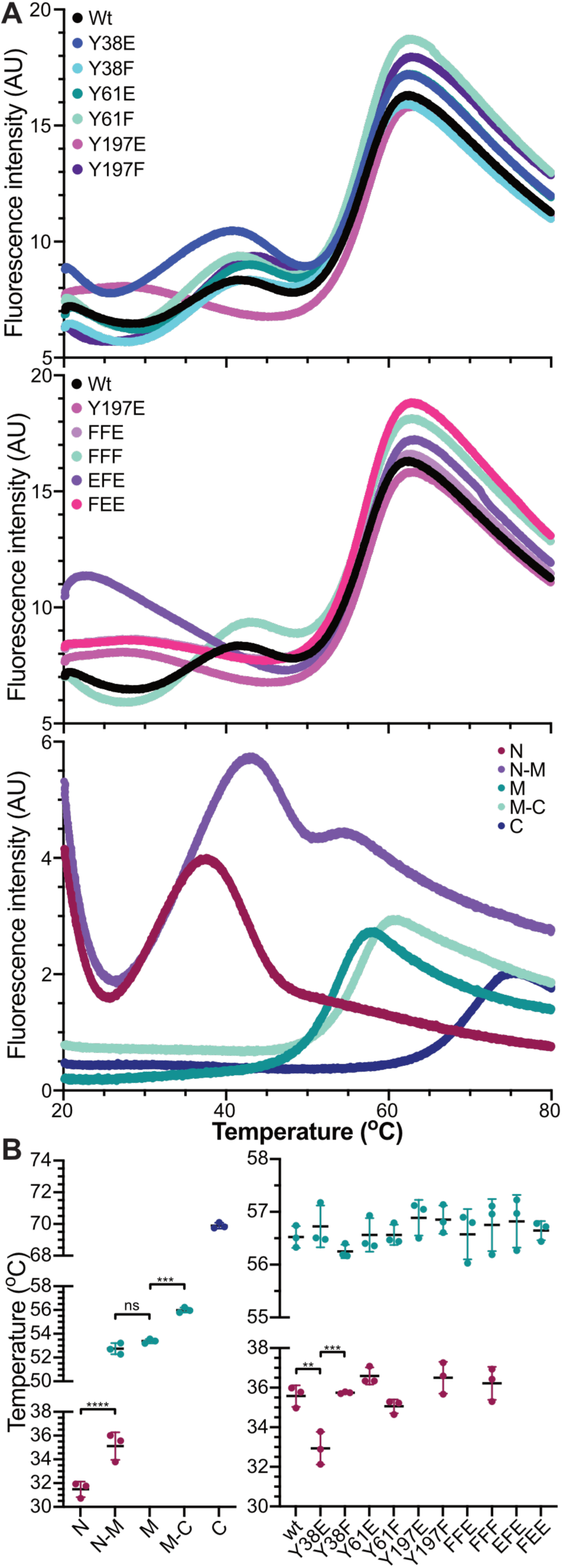
Some of the phosphomimetic amino acid replacements in hHsp90α significantly affect the conformation. **A**: Structural impact of the amino acid replacements in hHsp90α. Wild-type and mutant hHsp90α (upper two panels) and domain fragments of wild-type hHsp90α (lower panel) were analyzed by differential scanning fluorometry from 20 to 80°C using SYPRO orange. FFE: Y38F,Y61F,Y197E; FFF: Y38F,Y61F,Y197F; FEE: Y38F,Y61E,Y197E; EFE: Y38E,Y61F,Y197E; N, N-terminal nucleotide binding domain (amino acid residues 1-223); N-M, N-terminal and middle domain (1-545); M, middle domain (292-545); M-C, middle and C-terminal domain (292-732); C, C-terminal domain (545-732). **B**: Transition temperature calculated from the curves shown in A and similar curves by fitting the equation for thermal unfolding to each transition segment separately. Left panel, temperature transitions for the different domain constructs of wild-type hHsp90α. Right panel, temperature transitions for full-length wild-type and mutant hHsp90α variants. For Y197E containing variants the first transition is missing or below 20°C. Shown are mean and standard deviation. Statistical significance was tested with ordinary one-way ANOVA and Šidák’s multiple comparison; ns, not significant; *, p < 0.05; **, p < 0.01; ***, p < 0.001; ****, p < 0.0001.

Transition temperatures were determined by fitting the thermodynamic unfolding equation to each segment with fluorescence increase (**Fig. 5B**). For wild-type Hsp90α we observed two temperature transitions, one at 35.6 ± 0.5°C and the second at 56.5 ± 0.2°C. The first transition corresponded to local conformational changes in the N-domain (single transition of N 31.5 ± 0.6°C; first transition of N-M 35.1 ± 1.2°C) and the second transition to the conformational change of the middle domain (second transition of N-M 52.7 ± 0.5°C; single transition of M 53.4 ± 0.2°C; single transition of M-C 56.0 ± 0.2°C). Most of the Hsp90 variants also exhibited both transitions with the first one between 35.1 and 36.6°C and the second between 56.3 and 56.8°C. Significant differences were only observed for Y38E with a first transition at 32.9 ± 0.8°C and for all variants that contained Y197E, for which the first transition seemed to be below 20°C and could therefore not be determined. No differences were observed for the second transition of any of the Hsp90 variants analyzed in this study.

### The Hsp90 variants differ in their affinity for cochaperones

To chaperone clients like SHRs and kinases, Hsp90 cooperates with a number of cochaperones including Hop/Sti1 and p23/Sba1 for SHRs and Cdc37 and Aha1 for kinases. We, therefore, determined the affinity of wild-type and mutant Hsp90α to these cochaperones. We introduced an N- or C-terminal tetra-cysteine tag and labeled the proteins with FlAsH™-EDT2 to measure the interaction by fluorescence anisotropy (**Fig. S2**). To make sure the tag and the FlAsH label does not significantly affect the affinity of Hsp90α to the cochaperone, we verified that the labeled cochaperone can be displaced by the corresponding unlabeled protein.

The affinity of Hsp90α to Aha1 was only affected by glutamate in positions 38 and 197 for which we determined a significantly lower KD than for their corresponding phenylalanine replacement variants (**Fig. 6A**). The affinity of all other Hsp90 variants for Aha1 was within experimental error identical to the affinity of wild-type Hsp90α.

**Figure 6:**
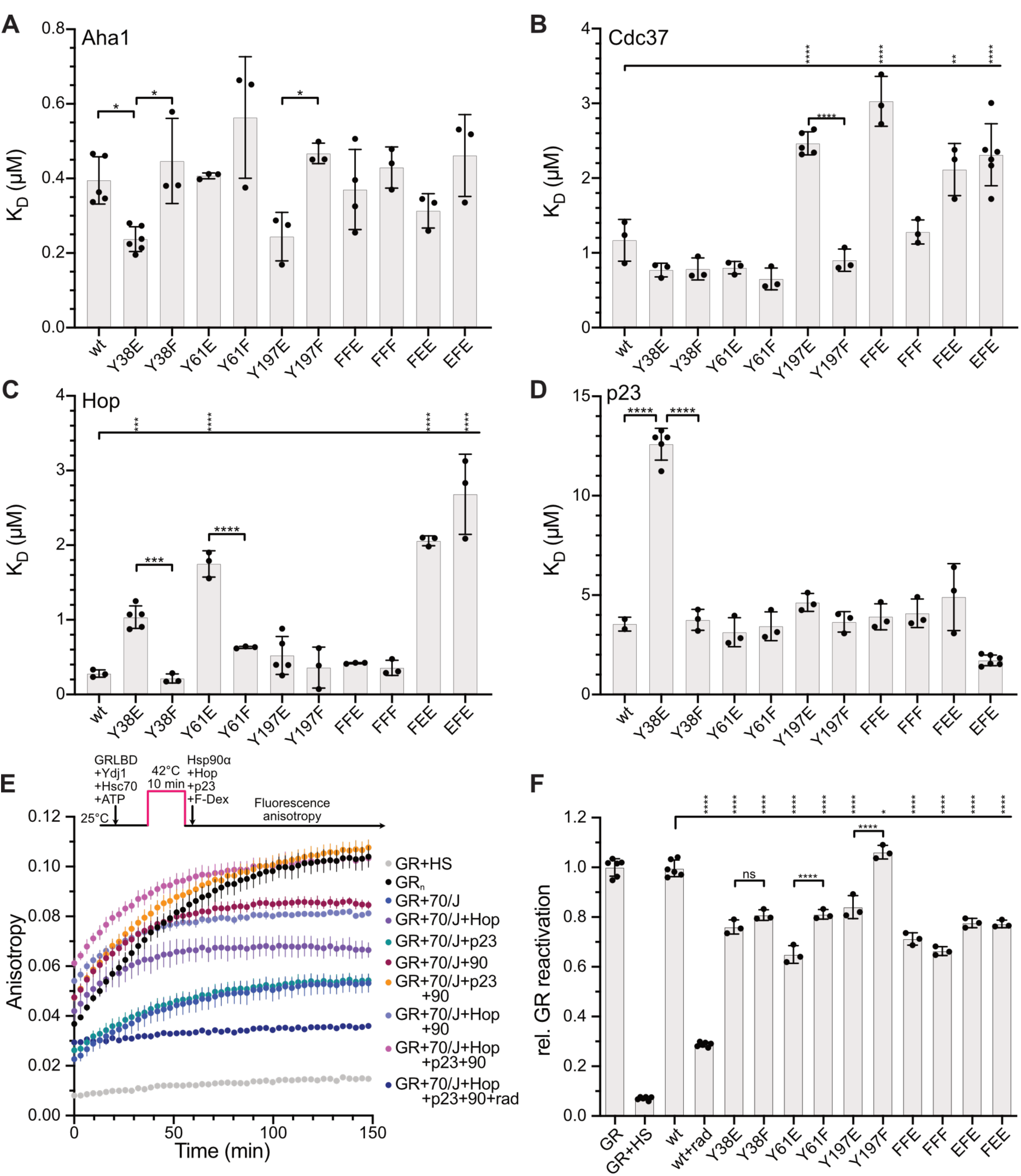
Phosphomimetic amino acid replacements in hHsp90α influence the binding of cochaperones and affect maturation of GRLBD *in vitro*. **A–D**: Dissociation equilibrium constants for the interaction of wild-type and mutant hHsp90α with N-terminally FlAsH-labeled Aha1 (A), N-terminally FlAsH-labeled CDC37 (B), N-terminally FlAsH-labeled Hop (C) and C-terminally FlAsH-labeled p23 (D) at room temperature. FFE: Y38F,Y61F,Y197E; FFF: Y38F,Y61F,Y197F; FEE: Y38F,Y61E,Y197E; EFE: Y38E,Y61F,Y197E. Shown are mean and standard deviation. Statistical significance was tested with ordinary one-way ANOVA and Šidák’s multiple comparison; *, p < 0.05; **, p < 0.01; ***, p < 0.001; ****, p < 0.0001. **E**: Refolding of denatured glucocorticoid receptor ligand binding domain (GR) by hHsp90α wild-type and mutant proteins in the presence of Hsc70, Ydj1, Hop, and p23. GR was denatured by incubation at 42°C for 10 min (HS) in the presence of Hsc70 and Ydj1 and reactivated by addition of hHsp90, Hop, and p23 at 25°C. GR reactivation was monitored by binding of fluorescein dexamethasone using fluorescence anisotropy. 70, Hsc70; J, Ydj1; 90, hHsp90α; rad, radicicol. For comparison, F-Dex binding to non-denatured, native glucocorticoid receptor ligand binding domain (GRn) is shown. **F**: Endpoint anisotropy values of reactions shown in E and similar reactions with mutant hHsp90α and all cochaperones. Shown are mean and standard deviation. Statistical significance was tested with ordinary one-way ANOVA and Šidák’s multiple comparison; ns, not significant; *, p < 0.05; **, p < 0.01; ***, p < 0.001; ****, p < 0.0001.

For the interaction of Hsp90 with Cdc37 only glutamate in position 197 had a significant impact (**Fig. 6B**). All Hsp90 variants that contained Y197E had a twofold increased KD for interaction with Cdc37 as compared to all other Hsp90 variants.

The interaction of Hsp90 with Hop was specifically affected by the glutamate replacement in positions 38 and 61 resulting in a 3.7 and 6.2-fold increased KD (**Fig. 6C**). This increase was even more pronounced when Y38E or Y61E were combined with Y197E. Replacement of only Y197 of Hsp90α with glutamate did not affect the interaction with Hop. The replacement of any of the investigated tyrosines with phenylalanine did not result in any significant impact on the affinity to Hop.

Since p23 requires a closed conformation of Hsp90 to bind [5], Hsp90 was incubated with AMPPNP, before mixing with p23 to induce a closed conformation. Despite this pre-incubation, it took at least 2 h to reach the binding equilibrium suggesting that in the presence of AMPPNP Hsp90α is in equilibrium between closed and open conformation until p23 stabilizes the closed conformation (**Fig. S2E & F**). For the interaction of Hsp90α with p23 only the replacement of Y38 with glutamate had a noticeable impact, increasing the KD 3.5-fold (**Fig. 6D, Fig. S2D**). This increase was canceled out by simultaneous replacement of Y197 with glutamate. Replacements of Y61 with glutamate or any of the three tyrosines with phenylalanine had not influence on the interaction with p23.

### Hsp90 variants differ in their ability to assist GR maturation in vitro

Glucocorticoid Receptor (GR) is one of the best-studied clients of Hsp90. Hsp90 binds to the C-terminal Ligand Binding Domain (LBD) of GR. Hsc70 with the J-domain cochaperone Ydj1 in the presence of ATP binds to GRLBD and unfolds the helix that forms the lid of the hormone-binding pocket, releasing the hormone in the process. Hop stabilizes the intermediate Hsc70-Hsp90-GRLBD complex and thereby mediates the transfer of unfolded GRLBD from Hsc70 to Hsp90. Cochaperone p23 facilitates the maturation of GRLBD by stabilizing the Hsp90-GRLBD complex [18, 19, 43, 44]. The matured GRLBD binds to the fluorescently labeled hormone Dexamethasone Fluorescein (F-Dex) and an increase in fluorescence anisotropy is observed. To increase the unfolding of GRLBD before binding to Hsp90, we subjected GRLBD to a heat shock (HS) treatment (42°C, 10 min) in the presence of Hsc70/Ydj1/ATP as described in [45].

Without heat shock and without any chaperones, native GRLBD bound F-Dex slowly and fluorescence anisotropy reached maximal values after ∼150 min. In contrast, GRLBD heat shocked without chaperones lost F-Dex binding ability almost completely (**Fig. 6E**). GRLBD heat-denatured in the presence of Hsc70/Ydj1 and subsequently incubated at room temperature regained some F-Dex binding ability and fluorescence anisotropy reached about 43% of the values observed with undenatured GRLBD. Addition of Hop, p23, or Hsp90 individually to GRLBD, heat denatured in the presence of Hsc70/Ydj1, increased the recovery of F-Dex binding ability not (p23) or only slightly (Hop and Hsp90) and only the combined addition of Hop, p23 and Hsp90 led to full recovery of F-Dex binding ability of GRLBD consistent with published data [44, 46]. In the presence of the Hsp90 inhibitor radicicol no time-dependent increase in fluorescence anisotropy was observed, indicating that the major part of GRLBD was not refolded. Replacing wild-type Hsp90α by the glutamate or phenylalanine variants reduced the recovery of F-Dex binding ability of GRLBD significantly except for Y197F, which was as active as wild-type Hsp90α (**Fig. 6F**). The lowest recovery of F-Dex binding ability was observed for Y61E.

## Discussion

In this study we investigated three previously by mass spectrometry identified tyrosine phosphorylation sites (Y38, Y61, Y197) in human Hsp90α *in vivo*, using a yeast model system and HepG2 HSP90AA1^−/−^ cells, and *in vitro*, using purified proteins. Our results are summarized in **Figure 7** and indicate that all three tyrosines are sensitive to replacement by glutamate and phenylalanine. Different clients were affected differently by the amino acid replacements, suggesting that phosphorylation of these tyrosines modulates the activity of Hsp90 towards individual clients distinctly. Although we could not test the effect of Y38E and Y61E on clients in yeast as these variants did not complement yeast growth, in human HepG2 cells at least Y61E stabilized two Hsp90α clients, DNA-PKcs and NBN, significantly better than the wild-type Hsp90α and the phenylalanine counterparts, indicating that a negative charge in this position is not always detrimental. Also, Y197E complemented yeast growth better than wild-type Hsp90 suggesting that some clients that are limiting for yeast growth are preferentially chaperoned by this variant. The molecular reason for this differential chaperoning of clients seems to be the alteration of the affinity of Hsp90 to cochaperones (**Fig. 7B**) and conformational changes as suggested by differential scanning fluorimetry. These data indicate that the investigated tyrosines are hotspots for modulation of Hsp90α‘s chaperone activity (**Fig. 7A**) and we propose that our data support the hypothesis that PTMs bias Hsp90’s activity towards certain clients that are preferentially chaperoned, while other clients are chaperoned less efficiently. This biasing of Hsp90 activity by posttranslational modification could be called Hsp90 code in analogy to Histone code.

**Figure 7:**
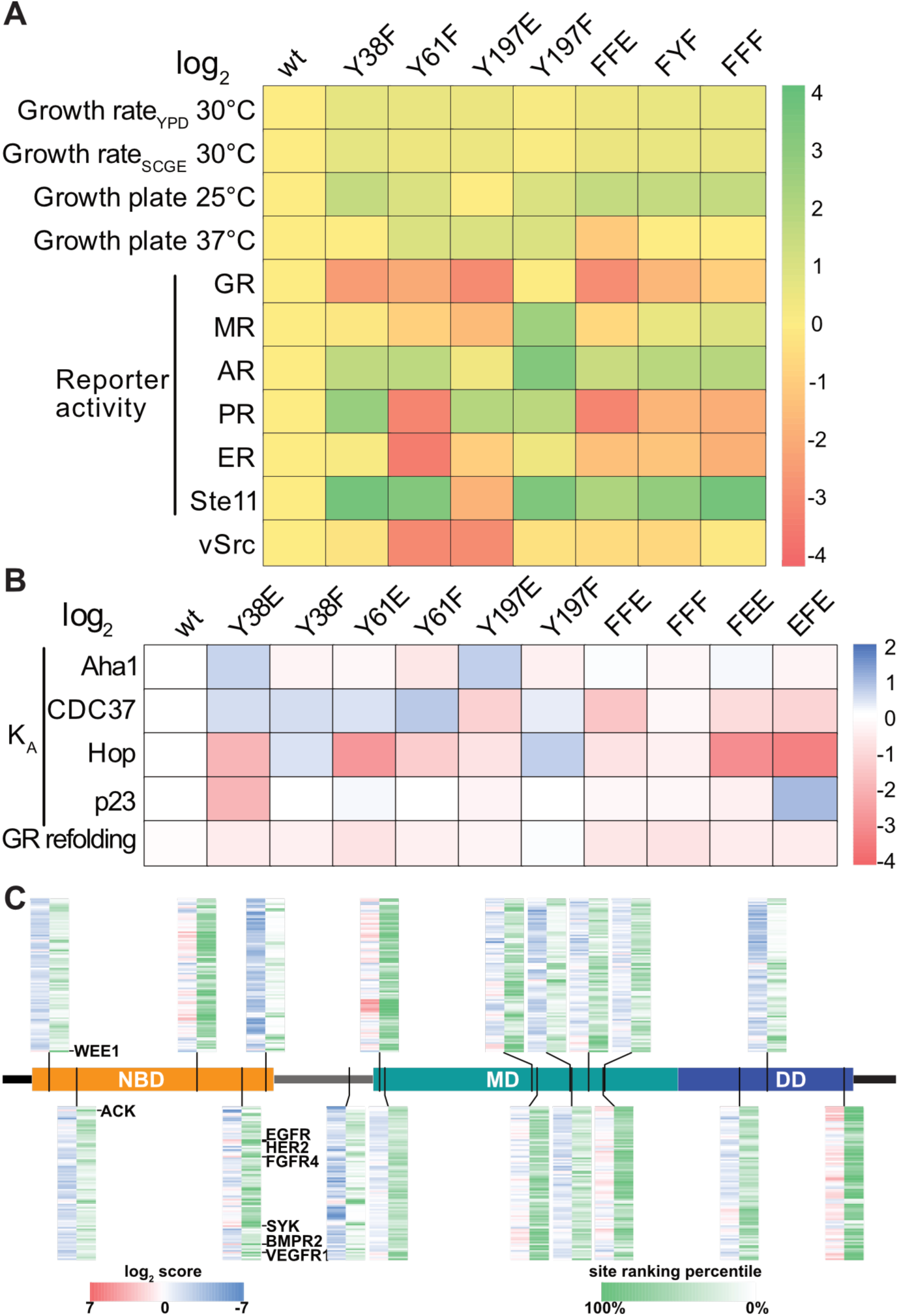
*In vivo* and *in vitro* data suggest that the three investigated tyrosine residues are switch points that tune hHsp90α for specific clients. **A**: Summary of the *in-vivo*-data in the yeast model system. Heat map of log2 transformed values for mutant hHsp90α relative to the corresponding value for wild-type hHsp90α. Color scheme is shown to the right. Growth rates in liquid YPD and SCGE medium at 30°C (data from Fig. 1E **and F**); growth on plates at 25°C and 37°C was estimated from the colony size (data from Fig. 1D); GR, MR, AR, PR, ER and Ste11, relative reporter expression measured as β-Galactosidase activity (data from Fig. 2); v-Src activity was quantified by calculating the relative increase in time a strain expressing V-SRC needed to reach an OD600 of 1 relative to the same strain containing the empty vector grown in parallel under the same conditions (data from Fig. 3C):
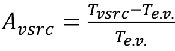; with T being the time to reach OD600 of 1. **B**: Summary of the *in-vitro*-data with purified proteins. Heat map of log2 transformed values for mutant hHsp90α relative to the corresponding value for wild-type hHsp90α. Association equilibrium constant KA (=1/KD) for the interaction of the mutant hHsp90α with Aha1, Cdc37, Hop, and p23 (data from **Figs. 6A-D**). Log2 transformed refolding yields for wild-type and mutant Hsp90α (fluorescence anisotropy values relative to native GRLBD; date from Fig. 6F). **C**: Each of the 18 tyrosines known to be phosphorylated is predicted to be phosphorylated by a unique set of tyrosine kinases. Domain structure of human Hsp90α with the indicated positions of the 18 tyrosines. For each site the tyrosine kinase prediction tool was used to generate score and percentile distribution for each of the 93 known tyrosine kinease in human cells (https://www.phosphosite.org/kinaseLibraryAction; not considering priming phosphorylation) [56, 63]. The result table was sorted in alphabetical order first according to kinase family and second for the individual kinase. Thus, the line that corresponds to a specific kinase is always at the same position within the heat map. The score (left; red to blue) indicates the log2 transformed preference of the given sequence over a random sequence containing the tyrosine in the middle. The percentile (right; green to white) indicates for each kinase the relative preference within the 5431 known phosphorylated tyrosine residues within the human proteome.

Our study has clear limitations. We did not investigate the effect of phosphorylation *per se* but of the replacement of tyrosine with glutamate, which seems to be a reasonably good surrogate for phosphorylated serine but mimics phosphorylated tyrosine poorly from a structural point of view. Although the non-natural amino acid 4-carboxy-L-phenylalanine might have been a better surrogate for phosphorylated tyrosine [47], the reduced expression levels associated with the nonsense suppression precluded its use in our study as human Hsp90α was already expressed to low levels and barely complemented the lethal phenotype of the lack of endogenous Hsp90 in yeast (see **Fig. 1E & F**). We have tested a considerable number of expression systems without achieving better expression levels and more robust complementation. Furthermore, we wanted to combine phosphomimetic variants with each other, which would imply the use of multiple stop codons within the same coding sequence when using non-natural amino acids and would reduce the expression level further, precluding comparison with wild-type and single mutants. From the three positions within Hsp90α we intended to analyze *in vivo* in this study, only Y197E complemented growth in yeast, whereas any variant that contained Y38E or Y61E did not complement. These results prompt the question whether glutamate in positions 38 and 61 compromise the structure of Hsp90α. Our DSF experiments showed thermal transition for Y61E that were within experimental error identical to wild-type Hsp90 (**Fig. 5A & B**), suggesting no structural impairment. For Y38E we found a first temperature transition at 32.9°C that was 2.6°C lower than the transition for wild-type Hsp90, suggesting some conformational changes in the nucleotide binding domain (**Fig. 5B**). For Y197E the first temperature transition seemed to be below 20°C, precluding exact determination with this method (**Fig. 5A**). Nevertheless, Y197E seemed to be functional *in vivo* as it complemented yeast growth even better than wild-type Hsp90α and Y197F (**Fig. 1E & F**), suggesting that this first transition does not represent a detrimental unfolding of the nucleotide binding domain. Y38E had a higher affinity for Aha1 than Y38F and wild-type Hsp90 *in vitro*, consistent with a published study in which human Hsp90α and its Y38F variant were immunoprecipitated from Cos7 cells coprecipitating Aha1 with the wild type but not with Y38F [26]. More importantly, yeast Hsp90 coprecipitated Aha1 from wild-type cells but not from cells in which swe1, the only tyrosine kinase in yeast, was deleted, whereas yHsp90-Y24F (homologous to Y38F in human Hsp90α) did not coprecipitate Aha1 from wild-type cells [26]. For CDC37 we did not observe any difference between Y38E, Y38F, and wild-type Hsp90 likewise consistent with published immunoprecipitation data [26]. In contrast, our interaction data for Hop and p23 binding are not consistent with the immunoprecipitation data in that study. Whereas we found a reduced affinity of Y38E for Hop and p23 as compared to wild type and Y38F (**Fig. 6C & D**), similar amounts of Hop and even increased amounts of p23 were precipitated with wild-type Hsp90α as compared to the amount coprecipitated with Y38F [26]. Such discrepancies might be due to further modifications that can occur in cells but were not present in our purified proteins. For Y61E the affinity to Hop was also markedly reduced. This observation is consistent with hydrogen exchange experiment with yeast Hsp90 and Sti1 (Hop homologue in yeast). When Sti1 interacted with yHsp90, a large protection was found in segment 43-62, which contains Y47 corresponding to Y61 in human Hsp90α [48]. This protection was even increased when TPR1 of yeast Sti1 was deleted. The Hsp70-Hop-Hsp90-GR loading complex shows Hop positioned in a way that could easily be imagined to interact with the region around Y61 in the absence of Hsp70 and GR [18]. Such an interaction might be disturbed by glutamate in position 61. For Y197E we found a reduced affinity for Cdc37 consistent with published data showing that Hsp90 released from a complex with Cdc37 is phosphorylated in Y197 [11], suggesting that Y197E recapitulates the behavior of phosphorylated Y197. Therefore, in our experiments glutamate seemed to be a suitable surrogate for phosphorylated tyrosine.

From a structural point of view the non-phosphorylatable phenylalanine seems to be very similar to the unphosphorylated tyrosine. However, the phenylalanine is much more hydrophobic with a hydrophathy index on the Kyte-Doolittle scale of 2.8, ranked between cysteine/cystine and leucine, whereas tyrosine has a hydrophathy index of −1.3, ranked between serine and proline [49]. Furthermore, tyrosine can engage in hydrogen bonds as donor as well as acceptor. We therefore examined several structures of human Hsp90α and generated homology models of human Hsp90α to different Hsp90 structures available using SWISS-MODEL [50, 51]. In many structures the phenolic hydroxyl group of Y197 formed hydrogen bonds, whereas Y38 and Y61 are involved in hydrogen bonding only in few models. Based on these observations, one could expect reduced stability and potentially a loss of function when replacing tyrosine with phenylalanine in these positions. However, all phenylalanine variants had temperature transitions that were within experimental error indistinguishable from the temperature transitions of wild-type Hsp90α (**Fig. 5B**), suggesting that they had no folding defects detectable by DSF. Likewise, none of the phenylalanine variants had KD-values significantly different from wild-type Hsp90α for the interaction with any of the four cochaperones tested in this study (**Fig. 6A-D**). This suggests that phenylalanine is a suitable surrogate for non-phosphorylated tyrosine at the investigated positions of Hsp90α in the context of interaction with these cochaperones. In contrast, Y38F and Y61F complemented yeast growth much better than wild-type human Hsp90α especially at low temperatures (**Fig. 1D - F**), suggesting a gain of function. As we could not detect phosphorylation of wild-type Y38 or Y61 by mass spectrometry, we consider the inactivation of a significant subpopulation of wild-type Hsp90 by phosphorylation of Y38 or Y61 by Swe1 kinase as explanation for the growth difference as unlikely. It was shown previously that amino acid replacements in the N-terminal domain of human Hsp90α that accelerate N-terminal docking and reopening support yeast growth better than wild-type Hsp90α [52]. The increased hydrophobicity of phenylalanine in positions 38 and 61 might therefore accelerate N-terminal docking of Hsp90, consistent with crosslinking data for yeast Hsp90-Y24F (corresponding to human Hsp90α-Y38F) [26].

Both Y38F and Y61F assisted maturation of glucocorticoid receptor much less efficient than wild-type Hsp90α and led to a roughly 80% reduction in reporter activity (**Fig. 2A**). The structure of the Hsp70-Hop-Hsp90-GRLBD loading complex provides a possible explanation [18]. In both Hsp90 protomers the hydroxyl group of Y61 forms polar contacts to the two Hsp70 molecules (D160) in the complex and thereby might stabilize the loading complex. Consistent with this hypothesis is also that *in vitro* Y61E supported GR reactivation significantly less well than Y61F (**Fig. 6F**). These results suggest that the negative charge of glutamate in position 61, leading to electrostatic repulsion of the aspartate in position 160 of Hsp70, would be more unfavorable for complex formation than the missing hydrogen bond in the Y61F variant. The hydroxyl group of Y38 in one of the two Hsp90 protomers, the one that contacts the free Hsp70 and Hop, forms internal hydrogen bonds to N40 and together with backbone contacts to T36 and K41 might stabilize a specific conformation of the N-terminal domain of that Hsp90 protomer. These hydrogen bonds are absent in the second protomer. Whether these internal contacts are sufficient to explain the reduced ability of Y38F to mature GR is not clear. The intermediate Hsp90-FKBP52-GR complex and the mature Hsp90-p23-GR complex show for neither Y38 nor Y61 any hydrogen bonds of the phenolic hydroxyl group [19, 53]. Y197F assists maturation of GR like wild-type Hsp90, though in both protomers of the loading complex Y197 forms internal hydrogen bonds with R202. These hydrogen bonds do not seem to be critical for GR maturation.

GR maturation by the Hsp70-Hsp90 machinery is considered to be a paradigm for the maturation of all steroid hormone receptors. Our data reveal similarities but also suggest that there might be specific differences consistent with earlier work on the cochaperone requirement of the different steroid hormone receptors [35]. Like for GR, Y61F could not assist activation of MR, PR and ER. These results suggest that Hsp70 interacts with Hsp90 during loading of these hormone receptors as suggested by the cryoEM structure of the Hsp70-Hsp90-Hop-GR loading complex [18], involving polar interactions with Y61. Although, Sti1/Hop does not seem to be essential for maturation of MR, AR, PR, and ER per se [35]. On the other side, MR and AR are much better chaperoned by Y197F than by wild-type Hsp90, whereas PR is chaperoned more efficiently by Y38F, contrasting the situation for GR (**Fig. 2**). These data hint that the loading conformation of Hsp90 could be different for MR, AR and PR as compared to the conformation in the GR loading complex.

Distinct Hsp90 requirements were also found for protein kinases. The serine/threonine kinase Ste11 dependent transcriptional reporter was induced only marginally in the presence of wild-type human Hsp90α but by an order of magnitude stronger by Y38F, Y61F, Y197F and the FFF Hsp90 variant (**Fig. 2F**). In the presence of Y197E barely any Ste11/Ste12 reporter activity was measured, consistent with the observation that all Y197E containing variants had a reduced affinity for the kinase specific cochaperone Cdc37 (**Fig. 6B**). These results are consistent with the published data that phosphorylation of Y197 by Yes kinase leads to the release of Cdc37 from the Hsp90-Cdc37-kinase client complex [11]. The tyrosine kinase v-Src was likewise not matured by Y197E, consistent with the reduced affinity to Cdc37 (**Fig. 3**). Interestingly, Y61F also did not mature v-Src (**Fig. 3B, C**), indicating that tyrosine kinases may have other requirements for Hsp90 than serine/threonine kinases. Surprisingly, the variant that combined the two amino acid replacements Y61F and Y197E that did not support v-Src maturation together with Y38F was able to recue v-Src activity, indicating that reduced interaction with Cdc37 and acceleration of different steps of the Hsp90 chaperone cycle could compensate each other. Taken together, we conclude that whether phenylalanine is a suitable surrogate for non-phosphorylated tyrosine depends on the position within Hsp90, the assay, the client and potentially also the cochaperones involved.

Interestingly, combining two favorable amino acid replacement in general did not result in synergistic or additive effects in supporting yeast growth or client maturation, whereas combining a favorable and an unfavorable amino acid replacement in general resulted in an intermediate phenotype, suggesting compensatory effects. If the single variants accelerated or decelerated the chaperone cycle at different stages, they would shift the conformational equilibria to stages that may be favorable for some clients but not for others. Multiple amino acid replacements might counteract each other shifting the conformational equilibria back to a more wild-type like situation.

Another limitation of our study, like any mutagenesis study, is that the dynamic and transient nature of PTMs cannot be mimicked. A phosphorylation event might be important for a specific stage of the chaperoning cycle but would arrest progression through the cycle, if not removed in time. Such a modification would appear negative in our study, whereas it might promote progression through the cycle and thereby allow more efficient maturation of the client in mammalian cells as suggested by a previous publication [11]. Therefore, we do not claim that phosphorylation of Y38, Y61 or Y197 *per se* is detrimental for chaperone activity of Hsp90 but that a permanent negative charge at these positions rendered Hsp90 inactive for some of the clients tested but not for others (compare [54]).

A surprising finding was that the differences in the ability of the Hsp90 variants to mature glucocorticoid receptor were much more pronounced in the yeast model system than in the *in-vitro*-GRLBD-refolding system where we observed for all variants significant refolding activity. There could by several explanations for these observations. (1) In the *in-vitro*-refolding assay the more stable F602S variant of the ligand binding domain of GR fused to maltose binding protein, similar to a published construct [16, 44] was used, whereas *in vivo* the full-length GR was present. (2) *In vitro* we have used the human cochaperones Hop and p23, whereas *in vivo* the yeast proteins Sti1 and Sba1 cooperated with human Hsp90α in the maturation of GR. The yeast proteins may have altered interaction kinetics with the different human Hsp90α variants. (3) Multiple cochaperones not related to GR maturation, most notably Cdc37, may compete with Sti1 and Sba1 for interaction with Hsp90, leaving limiting chaperoning capacity for GR. (4) *In vivo* Hsp90 has to take care of multiple clients at the same time while it chaperones GR. Such a competition may aggravate small differences in maturation efficiency. These differences between the yeast system as compared to our *in-vitro*-assays and potentially a mammalian *in vivo* system could also be considered advantageous as it seems to be more sensitive to small fitness differences of Hsp90, allowing to detect finetuning that might remain hidden in other systems.

Overall, our data suggest that the tyrosines in 38, 61 and 197 are potential regulators of client specificity consistent with previous cell biological [11, 26, 55] and structural data [18]. According to the recently mapped substrate specificity of all human kinases [56], the dual-specificity serine/threonine-tyrosine kinase Wee1, the negative regulator of G2-to-mitosis transition, is the top hit for phosphorylating Y38, consistent with earlier publications [26, 55]. For no other kinase the sequence motif containing Y38 was within the 90% percentile of substrates. In contrast, the kinase with the highest likelihood to phosphorylate Y61 is ACK1/TNK2, itself a client of Hsp90 and Cdc37 (Hsp90Int.DB; https://www.picard.ch/Hsp90Int/index.php) [41]. TNK2 is implicated in cell migration, cell survival, cell growth and proliferation. ACK1 regulates several Hsp90 clients including AKT1, EGFR, and androgen receptor [57, 58]. Phosphorylation of Hsp90 at Y61 might support this regulation. The sequence containing Y197 is in the >90% percentile of eleven kinases including BMPR2, SYK, MKK6/MAP2K6, HER2, EGFR, VEGFR1, FGFR4. Except for BMPR2, every single one of these tyrosine kinases interacts with Hsp90 or Cdc37 or both. Since the phosphomimetic Y197E has a lower affinity to the cochaperone Cdc37, which targets kinases to Hsp90, phosphorylation at Y197 might act as a negative feedback loop to prevent retargeting of Hsp90 to the kinase after an initial activation cycle, thereby contributing to limiting the activation of these signaling kinases. Thus, phosphorylation at each of the three sites, Y38, Y61, and Y197, by different kinases might bias Hsp90 to preferentially bind to some clients or to lower the ability to bind to other clients. In extension, many of the phosphorylation sites might have similar functional differences. Each of the 18 tyrosines, found by high-throughput mass spectrometry studies to be phosphorylated, has its own set of kinases as suggested by the different substrate prediction patterns (**Fig. 7C**). Some sites like Y38 and Y61 are predicted to be phosphorylated by only one or a few kinases (few dark red and dark green bands), whereas other sites like Y197 are phosphorylated by many kinases. All of these observations are consistent with the hypothesis of an Hsp90 code.

How could a Hsp90 code work? As many kinases are clients of Hsp90, maturation and activation of these kinases could lead to phosphorylation of Hsp90 reducing the activity of Hsp90 towards that particular kinase or kinases in general, limiting the overall activation of signaling pathways in a negative feedback loop. Furthermore, not all Hsp90 clients are expressed in every cell and specific modifications of Hsp90 could optimize Hsp90 for the needs of the particular cell type. As a consequence, cancer cells might reprogram Hsp90 for their needs by altering the Hsp90 code. The Hsp90 code could also be the underlying mechanism for the so-called epichaperome, Hsp90 complexes found in cancer cells and neurons in neurodegenerative diseases that seem to be maladaptive, promoting disease progression [59, 60].

## Supporting information

Supplemental Infomation

## Acknowledgements

We thank Stephan Hennes for excellent technical assistance. We thank D. Picard for the Δhsp82 Δhsc82 yeast strain and a HSP90AA1 expressing yeast plasmid. K. Morano for the Ste12-reporter plasmid, and J. Buchner for providing the plasmids for steroid hormone receptors and yeast plasmids with LacZ under the control of steroid response elements.

## Authors’ contributions

Conceptualization MPM; Formal analysis YH, RK, LL, MPM; Funding acquisition MPM, XC; Investigation YH, RK, LL; Methodology YH, RK, LL, RAK, MG, MPM; Project administration MPM, XC; Supervision MPM, XC; Visualization YH, RK, LL, MPM; Writing original draft YH, RK, MPM; Writing reviewing & editing YH, RAK, XC, MPM;

## References

[1] Biebl MM, Buchner J. Structure, Function, and Regulation of the Hsp90 Machinery. Cold Spring Harbor Perspectives in Biology. 2019;11:a034017–33.

[2] Taipale M, Krykbaeva I, Koeva M, Kayatekin C, Westover KD, Karras GI, et al. Quantitative analysis of HSP90-client interactions reveals principles of substrate recognition. Cell. 2012;150:987–1001.

[3] Miyata Y, Nakamoto H, Neckers L. The therapeutic target Hsp90 and cancer hallmarks. Current pharmaceutical design. 2013;19:347–65.

[4] Jarosz D. Hsp90: A Global Regulator of the Genotype-to-Phenotype Map in Cancers. Advances in cancer research. 2016;129:225–47.

[5] Ali MMU, Roe SM, Vaughan CK, Meyer P, Panaretou B, Piper PW, et al. Crystal structure of an Hsp90-nucleotide-p23/Sba1 closed chaperone complex. Nature. 2006;440:1013–7.

[6] Southworth DR, Agard DA. Species-dependent ensembles of conserved conformational states define the Hsp90 chaperone ATPase cycle. Molecular Cell. 2008;32:631–40.

[7] Mickler M, Hessling M, Ratzke C, Buchner J, Hugel T. The large conformational changes of Hsp90 are only weakly coupled to ATP hydrolysis. Nature Structural & Molecular Biology. 2009;16:281–6.

[8] Mader SL, Lopez A, Lawatscheck J, Luo Q, Rutz DA, Gamiz-Hernandez AP, et al. Conformational dynamics modulate the catalytic activity of the molecular chaperone Hsp90. Nature Communications. 2020;11:1410–12.

[9] Picard D. Chaperoning steroid hormone action. Trends in endocrinology and metabolism: TEM. 2006;17:229–35.

[10] Pratt WB, Toft DO. Steroid receptor interactions with heat shock protein and immunophilin chaperones. Endocrine reviews. 1997;18:306–60.

[11] Xu W, Mollapour M, Prodromou C, Wang S, Scroggins BT, Palchick Z, et al. Dynamic tyrosine phosphorylation modulates cycling of the HSP90-P50(CDC37)-AHA1 chaperone machine. Molecular Cell. 2012;47:434–43.

[12] Keramisanou D, Aboalroub A, Zhang Z, Liu W, Marshall D, Diviney A, et al. Molecular Mechanism of Protein Kinase Recognition and Sorting by the Hsp90 Kinome-Specific Cochaperone Cdc37. Molecular Cell. 2016;62:260–71.

[13] Meyer P, Prodromou C, Liao C, Hu B, Roe SM, Vaughan CK, et al. Structural basis for recruitment of the ATPase activator Aha1 to the Hsp90 chaperone machinery. The EMBO Journal. 2004;23:1402–10.

[14] Panaretou B, Siligardi G, Meyer P, Maloney A, Sullivan JK, Singh S, et al. Activation of the ATPase activity of hsp90 by the stress-regulated cochaperone aha1. Molecular Cell. 2002;10:1307–18.

[15] Hagn F, Lagleder S, Retzlaff M, Rohrberg J, Demmer O, Richter K, et al. Structural analysis of the interaction between Hsp90 and the tumor suppressor protein p53. Nature Structural & Molecular Biology. 2011;18:1086–93.

[16] Lorenz OR, Freiburger L, Rutz DA, Krause M, Zierer BK, Alvira S, et al. Modulation of the Hsp90 chaperone cycle by a stringent client protein. Molecular Cell. 2014;53:941–53.

[17] Karagöz GE, Duarte AMS, Akoury E, Ippel H, Biernat J, Morán Luengo T, et al. Hsp90-Tau complex reveals molecular basis for specificity in chaperone action. Cell. 2014;156:963–74.

[18] Wang RY, Noddings CM, Kirschke E, Myasnikov AG, Johnson JL, Agard DA. Structure of Hsp90-Hsp70-Hop-GR reveals the Hsp90 client-loading mechanism. Nature. 2022;601:460–4.

[19] Noddings CM, Wang RY, Johnson JL, Agard DA. Structure of Hsp90-p23-GR reveals the Hsp90 client-remodelling mechanism. Nature. 2022;601:465–9.

[20] Verba KA, Wang RY-R, Arakawa A, Liu Y, Shirouzu M, Yokoyama S, et al. Atomic structure of Hsp90-Cdc37-Cdk4 reveals that Hsp90 traps and stabilizes an unfolded kinase. Science. 2016;352:1542–7.

[21] Johnson JL. Mutations in Hsp90 Cochaperones Result in a Wide Variety of Human Disorders. Front Mol Biosci. 2021;8:787260.

[22] Johnson JL. Evolution and function of diverse Hsp90 homologs and cochaperone proteins. Biochimica et biophysica acta. 2012;1823:607–13.

[23] Schopf FH, Biebl MM, Buchner J. The HSP90 chaperone machinery. Nature reviews Molecular cell biology. 2017;18:345–60.

[24] Mollapour M, Tsutsumi S, Kim YS, Trepel J, Neckers L. Casein kinase 2 phosphorylation of Hsp90 threonine 22 modulates chaperone function and drug sensitivity. Oncotarget. 2011;2:407–17.

[25] Mollapour M, Tsutsumi S, Truman AW, Xu W, Vaughan CK, Beebe K, et al. Threonine 22 phosphorylation attenuates Hsp90 interaction with cochaperones and affects its chaperone activity. Molecular Cell. 2011;41:672–81.

[26] Mollapour M, Tsutsumi S, Donnelly AC, Beebe K, Tokita MJ, Lee M-J, et al. Swe1Wee1-dependent tyrosine phosphorylation of Hsp90 regulates distinct facets of chaperone function. Molecular Cell. 2010;37:333–43.

[27] Nguyen MTN, Knieß RA, Daturpalli S, Le Breton L, Ke X, Chen X, et al. Isoform-Specific Phosphorylation in Human Hsp90β Affects Interaction with Clients and the Cochaperone Cdc37. Journal of Molecular Biology. 2017;429:732–52.

[28] Bachman AB, Keramisanou D, Xu W, Beebe K, Moses MA, Vasantha Kumar MV, et al. Phosphorylation induced cochaperone unfolding promotes kinase recruitment and client class-specific Hsp90 phosphorylation. Nat Commun. 2018;9:265.

[29] Wang X, Lu XA, Song X, Zhuo W, Jia L, Jiang Y, et al. Thr90 phosphorylation of Hsp90alpha by protein kinase A regulates its chaperone machinery. Biochem J. 2012;441:387–97.

[30] Woodford MR, Truman AW, Dunn DM, Jensen SM, Cotran R, Bullard R, et al. Mps1 Mediated Phosphorylation of Hsp90 Confers Renal Cell Carcinoma Sensitivity and Selectivity to Hsp90 Inhibitors. Cell reports. 2016;14:872–84.

[31] Ogiso H, Kagi N, Matsumoto E, Nishimoto M, Arai R, Shirouzu M, et al. Phosphorylation analysis of 90 kDa heat shock protein within the cytosolic arylhydrocarbon receptor complex. Biochemistry. 2004;43:15510–9.

[32] Lu X-a, Wang X, Zhuo W, Jia L, Jiang Y, Fu Y, et al. The regulatory mechanism of a client kinase controlling its own release from Hsp90 chaperone machinery through phosphorylation. Biochemical Journal. 2014;457:171–83.

[33] Wider D, Peli-Gulli MP, Briand PA, Tatu U, Picard D. The complementation of yeast with human or Plasmodium falciparum Hsp90 confers differential inhibitor sensitivities. Mol Biochem Parasitol. 2009;164:147–52.

[34] Nordquist EB, English CA, Clerico EM, Sherman W, Gierasch LM, Chen J. Physics-based modeling provides predictive understanding of selectively promiscuous substrate binding by Hsp70 chaperones. PLoS Comput Biol. 2021;17:e1009567.

[35] Sahasrabudhe P, Rohrberg J, Biebl MM, Rutz DA, Buchner J. The Plasticity of the Hsp90 Co-chaperone System. Molecular Cell. 2017;67:947–61.e5.

[36] Liu L, Deng Y, Zheng Z, Deng Z, Zhang J, Li J, et al. Hsp90 Inhibitor STA9090 Sensitizes Hepatocellular Carcinoma to Hyperthermia-Induced DNA Damage by Suppressing DNA-PKcs Protein Stability and mRNA Transcription. Mol Cancer Ther. 2021;20:1880–92.

[37] Zhou X, Wen Y, Tian Y, He M, Ke X, Huang Z, et al. Heat Shock Protein 90α-Dependent B-Cell-2-Associated Transcription Factor 1 Promotes Hepatocellular Carcinoma Proliferation by Regulating MYC Proto-Oncogene c-MYC mRNA Stability. Hepatology (Baltimore, Md). 2019;69:1564–81.

[38] Beebe K, Mollapour M, Scroggins B, Prodromou C, Xu W, Tokita M, et al. Posttranslational modification and conformational state of Heat Shock Protein 90 differentially affect binding of chemically diverse small molecule inhibitors. Oncotarget. 2013;4:1065–74.

[39] Picard D, Khursheed B, Garabedian MJ, Fortin MG, Lindquist S, Yamamoto KR. Reduced levels of hsp90 compromise steroid receptor action in vivo. Nature. 1990;348:166–8.

[40] Xu Y, Lindquist S. Heat-shock protein hsp90 governs the activity of pp60v-src kinase. Proceedings of the National Academy of Sciences of the United States of America. 1993;90:7074–8.

[41] Echeverria PC, Bernthaler A, Dupuis P, Mayer B, Picard D. An interaction network predicted from public data as a discovery tool: application to the Hsp90 molecular chaperone machine. PLoS ONE. 2011;6:e26044.

[42] Louvion JF, Abbas-Terki T, Picard D. Hsp90 is required for pheromone signaling in yeast. Molecular biology of the cell. 1998;9:3071–83.

[43] Dittmar KD, Hutchison KA, Owens-Grillo JK, Pratt WB. Reconstitution of the steroid receptor.hsp90 heterocomplex assembly system of rabbit reticulocyte lysate. The Journal of biological chemistry. 1996;271:12833–9.

[44] Kirschke E, Goswami D, Southworth D, Griffin PR, Agard DA. Glucocorticoid receptor function regulated by coordinated action of the hsp90 and hsp70 chaperone cycles. Cell. 2014;157:1685–97.

[45] Moran Luengo T, Mayer MP, Rudiger SGD. The Hsp70-Hsp90 Chaperone Cascade in Protein Folding. Trends Cell Biol. 2019;29:164–77.

[46] Morishima Y, Kanelakis KC, Silverstein AM, Dittmar KD, Estrada L, Pratt WB. The Hsp organizer protein hop enhances the rate of but is not essential for glucocorticoid receptor folding by the multiprotein Hsp90-based chaperone system. The Journal of biological chemistry. 2000;275:6894–900.

[47] Xie J, Supekova L, Schultz PG. A Genetically Encoded Metabolically Stable Analogue of Phosphotyrosine in Escherichia coli. ACS chemical biology. 2007;2:474–8.

[48] Lee C-T, Graf C, Mayer FJ, Richter SM, Mayer MP. Dynamics of the regulation of Hsp90 by the co-chaperone Sti1. The EMBO Journal. 2012;31:1518–28.

[49] Kyte J, Doolittle RF. A simple method for displaying the hydropathic character of a protein. Journal of Molecular Biology. 1982;157:105–32.

[50] Waterhouse A, Bertoni M, Bienert S, Studer G, Tauriello G, Gumienny R, et al. SWISS-MODEL: homology modelling of protein structures and complexes. Nucleic Acids Res. 2018;46:W296–W303.

[51] Biasini M, Bienert S, Waterhouse A, Arnold K, Studer G, Schmidt T, et al. SWISS-MODEL: modelling protein tertiary and quaternary structure using evolutionary information. Nucleic Acids Research. 2014;42:W252–8.

[52] Reidy M, Masison DC. Mutations in the Hsp90 N Domain Identify a Site that Controls Dimer Opening and Expand Human Hsp90alpha Function in Yeast. J Mol Biol. 2020;432:4673–89.

[53] Noddings CM, Johnson JL, Agard DA. Cryo-EM reveals how Hsp90 and FKBP immunophilins co-regulate the glucocorticoid receptor. Nat Struct Mol Biol. 2023.

[54] Soroka J, Wandinger SK, Mäusbacher N, Schreiber T, Richter K, Daub H, et al. Conformational switching of the molecular chaperone Hsp90 via regulated phosphorylation. Molecular Cell. 2012;45:517–28.

[55] Mollapour M, Tsutsumi S, Neckers L. Hsp90 phosphorylation, We e1 and the cell cycle. Cell cycle (Georgetown, Tex). 2010;9:2310–6.

[56] Yaron-Barir TM, Joughin BA, Huntsman EM, Kerelsky A, Cizin DM, Cohen BM, et al. The intrinsic substrate specificity of the human tyrosine kinome. Nature. 2024;629:1174–81.

[57] Grovdal LM, Johannessen LE, Rodland MS, Madshus IH, Stang E. Dysregulation of Ack1 inhibits down-regulation of the EGF receptor. Exp Cell Res. 2008;314:1292–300.

[58] Mahajan K, Mahajan NP. Shepherding AKT and androgen receptor by Ack1 tyrosine kinase. J Cell Physiol. 2010;224:327–33.

[59] Rodina A, Xu C, Digwal CS, Joshi S, Patel Y, Santhaseela AR, et al. Systems-level analyses of protein-protein interaction network dysfunctions via epichaperomics identify cancer-specific mechanisms of stress adaptation. Nat Commun. 2023;14:3742.

[60] Chiosis G, Digwal CS, Trepel JB, Neckers L. Structural and functional complexity of HSP90 in cellular homeostasis and disease. Nat Rev Mol Cell Biol. 2023;24:797–815.

[61] Shiau AK, Harris SF, Southworth DR, Agard DA. Structural Analysis of E. coli hsp90 reveals dramatic nucleotide-dependent conformational rearrangements. Cell. 2006;127:329–40.

[62] Obermann WM, Sondermann H, Russo AA, Pavletich NP, Hartl FU. In vivo function of Hsp90 is dependent on ATP binding and ATP hydrolysis. Journal of Cell Biology. 1998;143:901–10.

[63] Johnson JL, Yaron TM, Huntsman EM, Kerelsky A, Song J, Regev A, et al. An atlas of substrate specificities for the human serine/threonine kinome. Nature. 2023;613:759–66.

