## Supplemental Infomation for "Modification of regulatory tyrosines biases human Hsp90α for interaction with cochaperones and clients"

### Supplemental Information

Supplemental materials and methods

Supplemental Figures S1 and S2

Supplemental Tables S1 and S2

#### Supplemental Materials and Methods

*Media and strains*

For non-selective growth, yeast was cultured in YPD (1% yeast extract, 2% peptone, 2% glucose, with 0.01% adenine). For selective growth, synthetic complete (SC) dextrose medium was used (SD, 2% glucose, 0.17% yeast nitrogen base without ammonium sulphate, 0.5% ammonium sulphate, with all amino acids except when indicated (dropout medium). For non-repressive growth or induction of the GAL1 promoter glucose was replaced by raffinose or galactose as indicated. SCGE medium, used for growth assays, is identical to SD medium except that it contains 3% glycerol and 2% ethanol instead of glucose. SD plates contained 1.5% agar (Becton Dickinson). For plasmid shuffling 10 mg/L 5'-fluoro-orotic-acid (5FOA, 100 mg/mL, Zymo Research Epigenetics Company) was added in dropout plates. All cells were grown at 30 °C unless otherwise stated.

*Spot assay*

For the spot assay overnight yeast cultures were first diluted to $OD_{600}$ = 0.2, from these diluted cultures serial 1:5 dilutions were prepared and 5 µl spotted onto the agar plate.

*Immunoblotting of yeast samples*

Yeast proteins were extracted according to a published protocol [1]. The yeast cells were pelleted by centrifugation (1000g, 5 min) and resuspended in water to $OD_{600}$ = 2.5. 1 volume of NaOH (0.2 M) was added and incubated at room temperature for 5 min before a second centrifugation (1000g, 5 min). The cells were resuspended in 1xSDS sample buffer 100 µL/5 OD (5% glycerol, 2% SDS, 4% β-mercaptoethanol, 0.06 M Tris-HCl pH 6.8, 0.0025%

bromophenol blue). The samples were heated at 88°C for 3 minutes and centrifuged to remove cell debris at maximum speed for 2 min. The supernatant was used for SDS-PAGE. The proteins were transferred to a polyvinylidenfluorid (PVDF) membrane with a semidry blotting system (BIO-RAD), and the membrane was blocked in 5% milk diluted in TBST (20 mM Tris-base, 150 mM NaCl, pH 7.6 (adjusted with HCl), 0.1% Tween 20 (v/v)). Incubation and imaging with antibodies were performed according to the manufacturer's instructions.

*Plasmid shuffling*

After transforming the parental strain (MPMH2458) with the plasmid containing wild-type or mutant HSP90AA1, the cells were first selected on SD -Leu/-Ura plates for 48 h. SD -Leu medium was inoculated with a single clone. After Culturing for 48 h, the spot assay was performed on SD -Leu plates and SD -Leu/5FOA plates. These plates were incubated at 30 °C for 96 h. Clones that could complement and grow on the SD -Leu/5FOA plates were confirmed by re-streaking onto SD -Leu/5FOA plate again. The replacement of yHsc82 by human Hsp90α was confirmed by immunoblotting.

*Yeast growth assay in liquid culture and growth curve*

Yeast media (YPD or SCGE) were inoculation with a single colony from a freshly streaked plate. After overnight shaking at 30°C, the cultures were diluted to $OD_{600}$ 0.2 and continuously incubated for another 4 h. Then each sample was diluted to $OD_{600}$ 0.05 in 1 mL of the indicated medium. The samples were incubated in 24-well plates in a plate reader (SPECTRO-star Nano, BMG LABTECH) at 30°C with continuous shaking at 600 rpm for 48 h. Growth monitored by measuring the $OD_{600}$ in each well every 10 min. The results were analyzed in GraphPad Prism, and the doubling times and the growth rate constants were calculated by fitting an exponential growth equation to the datapoints below $OD_{600}$ of 0.5.

*In vivo reporter assay*

Yeast strains containing wild-type or mutant hHsp90α were transformed with a lacZ reporter plasmid, as well as the hormone receptor encoding plasmids (as described above). The reporter assay was established for five steroid hormone receptors GR, ER, MR, AR and PR, and for the kinase Ste11. Inducers to activate the reporter were 11-Deoxycorticosteron (DOC) for GR, dihydrotestosterone (DHT) for AR, β-estradiol for ER, progesterone for PR, aldosterone for MR (all hormones from Merck, Darmstadt, Germany) and α-factor for pheromone response (Shanghai RoyoBiotech co. Ltd, purity > 99 %).

The transformed strains were grown overnight at 30°C in SD-media before being diluted into two cultures with $OD_{600}$ 0.5. One of the two cultures was induced with 10 μM inducer (as described above). Both induced and non-induced cultures were then incubated for 3 h at 30°C.

The culture $OD_{600}$ was then recorded and 200 – 400 µL of the culture were harvested by centrifugation (1000 g, 5 min). The cells were then lysed by re-suspension into 800 µL buffer Z (10.68 g/L $NaH_2PO_4 \cdot 2H_2O$, 0.75 g/L KCl, 0.246 g/L $MgSO_4 \cdot 7H_2O$, β-mercaptoethanol 2.7 µL/mL, adjusted to pH 7.0 with HCl), followed by adding 50 µL $CHCl_3$ and 50 µL 0.1% SDS. The reporter reaction was triggered by adding 160 µL ONPG (4 mg/mL, Roth) to the lysate and incubating at 30°C until the development of a strong yellow color for a maximum time of one hour. The reaction was quenched with 400 µL of 1 M $Na_2CO_3$. The cellular debris was removed by centrifugation in a microfuge (14000 rpm), and 200 µL of the supernatant of each sample were transferred to a 96-well plate. The absorbance at 420 nm was measured with the plate reader (SPECTRO-star Nano, BMG LABTECH). The reporter activity was calculated using the following equation:

$$\beta - gal\ units\ = \frac{OD_{420}}{t \cdot V \cdot OD_{600}} \cdot 1000$$

Where $OD_{420}$ = supernatant absorbance at 420nm; t = incubation time (min) at 30°C after adding ONPG; V = volume of culture (mL); $OD_{600}$ = OD of the culture.

*v-Src toxicity assay*

Yeast strains containing wild-type or mutant hHsp90α were transformed with a 2µ-episomal plasmid expressing *V-SRC* under the control of a GAL1 promoter or the empty vector as the negative control. The cells were first grown in SC raffinose/-His medium overnight before performing the spot assay on SC galactose/-His plates and monitoring the growth in SC galactose/-His medium.

*Protein expression*

Lysogeny broth (LB) (~200 mL) containing the appropriate antibiotic was inoculated with colonies from an LB agar plate with freshly transformed BL21(DE3) pCodonPlus cells and allowed to grow overnight at 37°C. 2L 2xYT media containing the appropriate antibiotics were inoculated with the pre-culture and grown at 37°C until an OD of 1. The temperature was lowered to 18°C for GRLBD and to 25°C for all other proteins, and the cells were allowed to acclimate for 30 min. Protein expression was induced by adding 0.5 mM IPTG, and expression was continued overnight. The next day, cells were harvested by centrifugation at 5000 rpm and 4°C for 10 min. Cells were collected and stored at -80°C until protein purification. For GRLBD, 150 µM dexamethasone was added to the culture along with IPTG.

*Protein Purification*

Cell pellets were resuspended in 5 ml lysis buffer (20 mM HEPES pH 7.5, 100 mM KCl, 5 mM $MgCl_2$, 10% glycerol, 2mM DTT) per g of cell pellet, supplemented with 10 µg/mL Aprotinin,

10 µg/mL Leupeptin, 8 µg/mL Pepstatin, 2 mM PMSF, 1 mg/mL DNase I, and lysed using a microfluidizer, followed by centrifugation at 13000 rpm for 30 min to remove cell debris.

For Hsp90 the cell lysate was incubated with Protino Ni-IDA (Macherey-Nagel) at 4°C and then added to a gravity flow column. The column was subsequently washed with 10 column volumes of lysis buffer (supplemented with 30 mM imidazole) and eluted with buffer containing 400 mM imidazole. Desalting to a low salt buffer (20 mM HEPES pH 7.5, 20 mM KCl, 5 mM $MgCl_2$, 0.5 mM EDTA, 10% glycerol) was performed on a HiPrep 26/10 desalting column (Cytiva) to remove the imidazole. The protein was further purified with a Resource™ Q 6 mL column (Cytiva) with a 15 CV gradient from 20 mM KCl to 1 M KCl. Fractions containing the desired protein were pooled and the SUMO tag cleaved by the SUMO protease Ulp1 overnight, dialyzed against the storage buffer (40 mM HEPES pH 7.5, 150 mM KCl, 5 mM $MgCl_2$, 15% glycerol, 2 mM DTT). The cleaved proteins were purified by size exclusion chromatography on a HiLoad Superdex 200™ column (Cytiva) in the storage buffer. Protein fractions were analyzed by SDS-PAGE, pooled according to purity, concentrated to ~50 µM, flash frozen in liquid nitrogen, and stored at -80°C.

For Aha1, the same protocol was used except a Resource™ Q 1 mL column (Cytiva) and a HiLoad Superdex 75™ column (Cytiva) were used with storage buffer (25 mM HEPES pH 7.5, 50 mM KCl, 5 mM $MgCl_2$, 10% glycerol, 2 mM DTT). The purified protein was stored at -80°C.

For Hop, p23, Cdc37, and Hsp90α fragments, following Ni-IDA purification and overnight SUMO cleavage, another round of Ni-IDA affinity purification was performed. The purified protein was stored in a storage buffer (25 mM HEPES pH 7.5, 150 mM KCl, 5 mM $MgCl_2$, 10% glycerol, 2 mM DTT) at -80°C.

The MBP-GRLBD-F602S fusion construct was used to improve the solubility of the protein. The cell lysate was incubated with amylose resin and added to a gravity column. The column was washed with 10 column volumes of the lysis buffer (25 mM HEPES pH 7.5, 100 mM KCl, 5 mM $MgCl_2$, 10% glycerol, 2 mM DTT) supplemented with 50µM dexamethasone and eluted with 50 mM maltose. The protein was further purified using size exclusion chromatography on a Superdex® 200 10/300 GL and dialyzed for 2 h in the presence of 50 µM dexamethasone into storage buffer (25 mM HEPES/KOH pH 7.5, 100 mM KCl, 5 mM $MgCl_2$, 10% glycerol, 2 mM DTT).

For Hsc70, the same protocol was followed as for Cdc37 until the first round of Ni-IDA affinity purification. After overnight dialysis in the presence of Ulp1 protease, the protein was incubated with fresh Protino Ni-IDA Resin for 25 min and added to the gravity flow column. The purest fractions of the flow-through as confirmed by SDS-PAGE were pooled and dialyzed into low salt buffer (25 mM HEPES/KOH pH 7.5, 10 mM KCl, 5 mM $MgCl_2$, 10% glycerol) and further purified using a 1-mL-Resource™ Q column (Cytiva) with a 15 CV gradient from 10 mM KCl to 1 M KCl. The purified protein was dialyzed against storage buffer (40 mM HEPES pH 7.5, 150 mM KCl, 5 mM $MgCl_2$, 10% glycerol, 2 mM DTT) and stored at-80°C.

For Ydj1, the same protocol was followed as for Cdc37 until the first round of Ni-IDA affinity purification. After overnight dialysis in the presence of Ulp1 protease, the protein was further purified by size exclusion chromatography on a Superdex® 75 10/300 GL column in SEC buffer (25 mM HEPES/KOH pH 7.5, 300 mM KCl, 5 mM $MgCl_2$, 10% glycerol, 2 mM DTT). The purified protein was dialyzed against storage buffer (40 mM HEPES pH 7.5, 150 mM KCl, 5 mM $MgCl_2$, 10% glycerol), flash frozen in liquid N2 and stored at -80°C.

*Binding Affinity Measurements*

All cochaperones had at their N- or C-terminus a tetra-cysteine tag (ACCPGCCSGG). Cochaperones and Hsp90 were thawed and centrifuged at 20,000 g for 30 min at 4°C to remove aggregates, followed by buffer exchange to the corresponding reaction buffer using Zeba™ Spin Desalting Columns (Thermo Fisher Scientific). The cochaperone was used at a concentration of 5 µM and incubated with 10 µM of FlAsH-$EDT_2$™ and 2 mM TCEP at 4°C for 2 h to complete labeling. The FlAsH-labeled cochaperones were desalted using a Zeba™ Spin Desalting Columns to remove the unbound dye. For the fluorescence anisotropy assay the following buffers were used for each cochaperone: Aha1 (20 mM HEPES/KOH pH 7.5, 5 mM $MgCl_2$, 2% glycerol, 2 mM TCEP); Cdc37 (25 mM HEPES/KOH pH 7.5, 150 mM KCl, 5 mM $MgCl_2$, 2% glycerol, 2 mM TCEP); Hop (25 mM HEPES/KOH pH 7.5, 100 mM KCl, 5 mM $MgCl_2$, 2% glycerol, 2 mM TCEP); p23 (25 mM HEPES/KOH pH 7.5, 20 mM KCl, 5 mM $MgCl_2$, 2% glycerol, 2 mM TCEP).

#### Supplemental figures

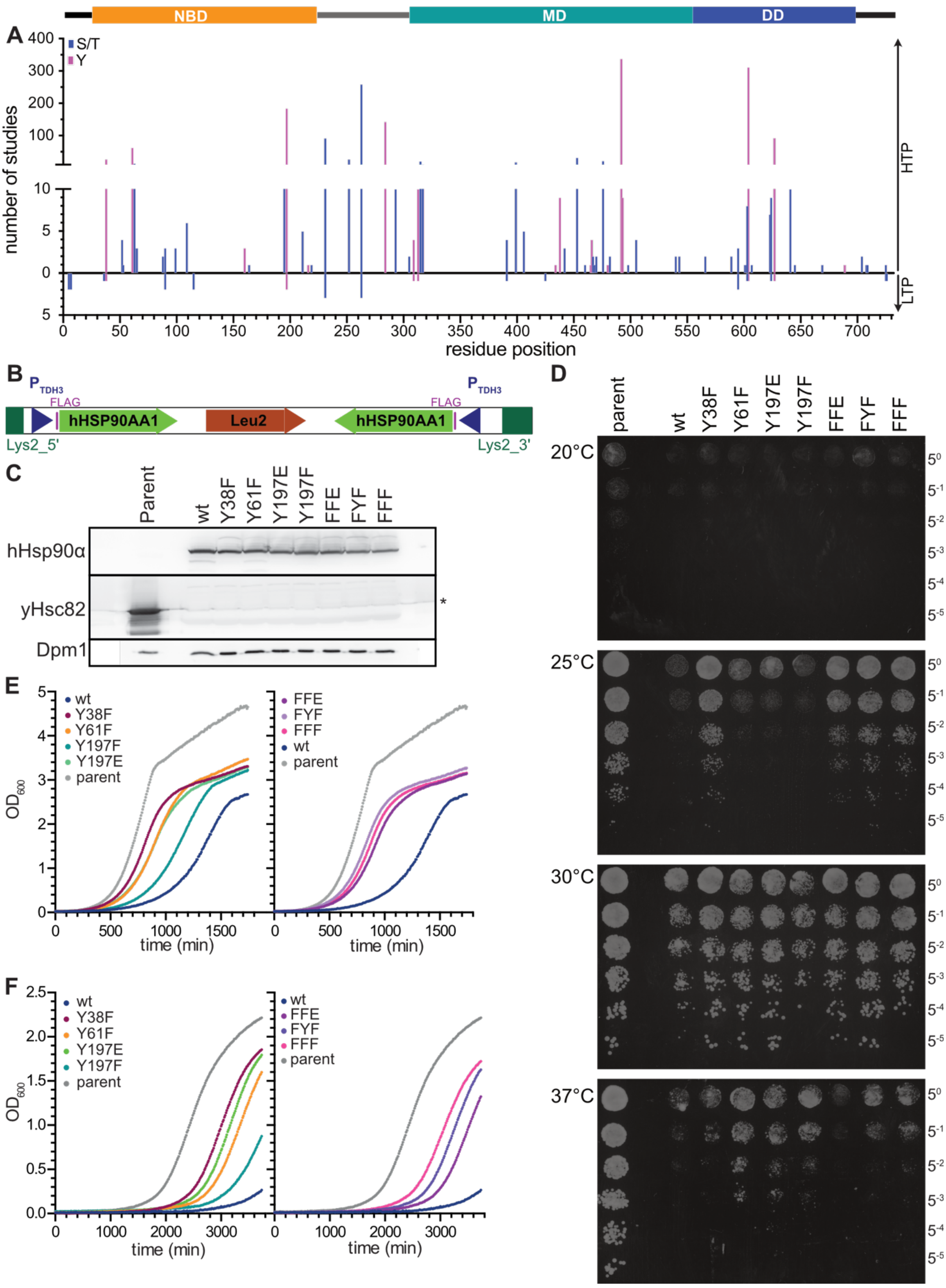

**Supplemental Figure S1 related to Figure 1: Phosphomimetic and non-phosphorylatable variants of human Hsp90α differentially support growth of *S. cerevisiae* cells lacking endogenous Hsp90.**

**A**, Number of high-throughput mass spectrometry studies (upwards) and low-throughput cell biological or biochemical studies (downwards) identifying the respective residue to be phosphorylated.

**B**, expression cassette for human *HSP90AA1* genes that was integrated into the LYS2 locus in the yeast strain MPMH2458.

**C**, Immunoblot identifying yHsc82 or hHsp90α in the strains after plasmid shuffling. The asterisk indicates an artifact that is even visible in lanes where no extracts were loaded.

**D**, Temperature sensitivity of yeast cells expressing wild-type or mutant human *HSP90AA1* as sole source for Hsp90. Fivefold dilutions of overnight cultures of the MPMH2458 strain expressing high levels of yHsc82 (parental) or wild-type or mutant hHsp90α were spotted on YPD plates and grown at the indicated temperatures for 48 h (growth after 72 h is shown in **Fig. 1D**).

**E & F**, data from Fig. 1E & F split into two panels for more clarity.

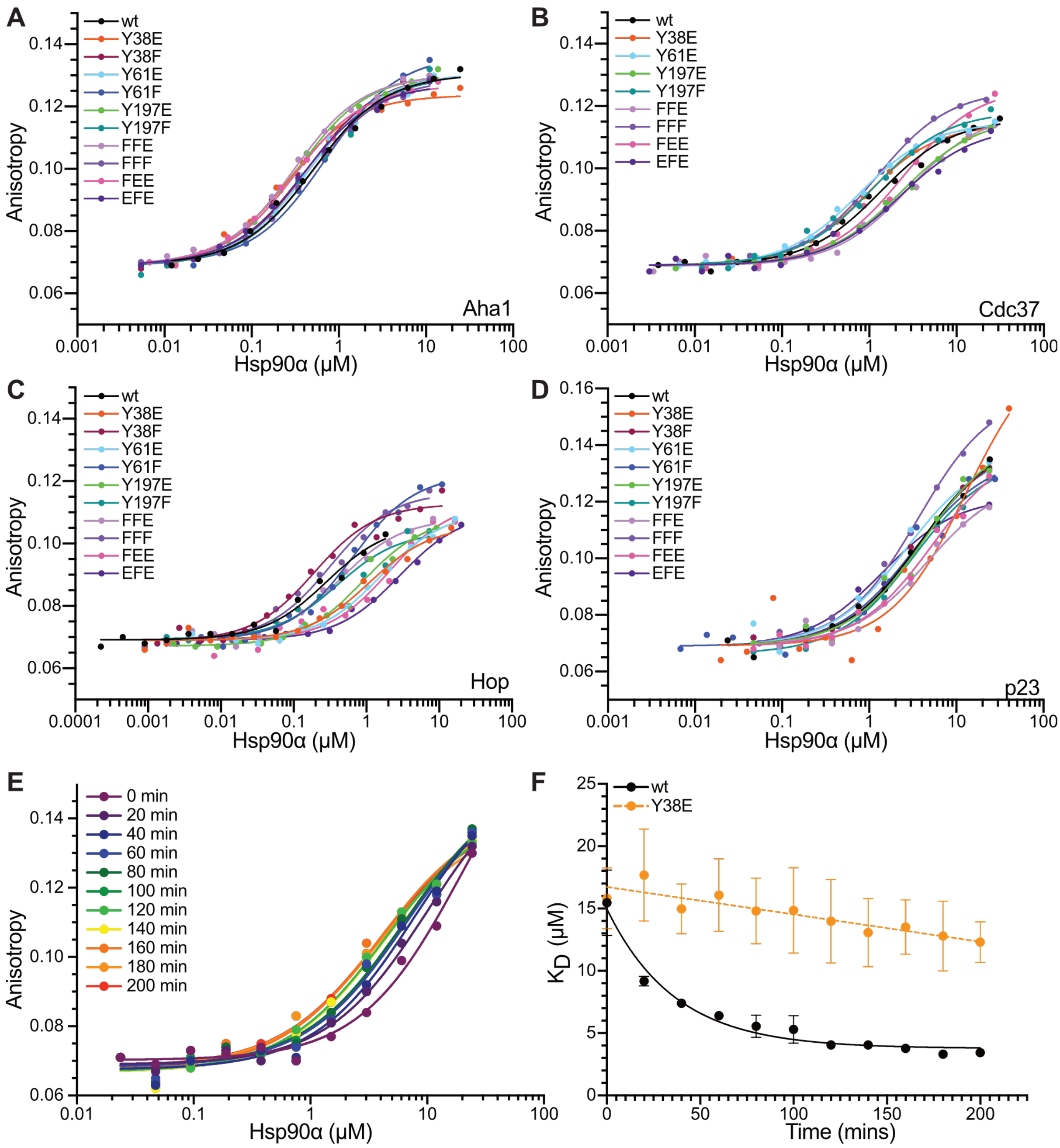


**Supplemental Figure S2 related to Figure 6: Phosphomimetic amino acid replacements in hHsp90α influence the binding of cochaperones**

**A-D**, Sample data of the dissociation equilibrium titration for the interaction of wild-type and mutant hHsp90α with N-terminally FlAsH-labeled Aha1 (**A**), N-terminally FlAsH-labeled Cdc37 (**B**), N-terminally FlAsH-labeled Hop (**C**) and C-terminally FlAsH-labeled p23 (endpoint) (**D**). FFE: Y38F,Y61F,Y197E; FFF: Y38F,Y61F,Y197F; FEE: Y38F,Y61E,Y197E; EFE: Y38E,Y61F,Y197E. Curves are fits of the quadratic solution of the law of mass action to the data.

**E & F**, Apparent binding affinity of Hsp90 to p23 was dependent on incubation time. Time dependence of titration curves (**E**) and calculated apparent $K_D$ values (**F**) for the interaction of wild-type hHsp90α (**E & F**) and Y38E (**F**) with p23. Error bars represent the standard deviation of the data.

**Table S1: Plasmids used in this study**

| Plasmid | Backbone | Insert | Reference |
|---|---|---|---|
| pLys2LR | pBluescript SK | *S. cerevisiae* Lys2 fragments, *K. lactis LEU2* | [2] |
| p1760 | pLys2LR | (TDH3-FLAG-hHsp90α-wt)x2 | This study |
| p1473 | pLys2LR | (TDH3-FLAG-hHsp90α-Y38E)x2 | This study |
| p1474 | pLys2LR | (TDH3-FLAG-hHsp90α-Y38F)x2 | This study |
| p1475 | pLys2LR | (TDH3-FLAG-hHsp90α-Y61E)x2 | This study |
| p1476 | pLys2LR | (TDH3-FLAG-hHsp90α-Y61F)x2 | This study |
| p1477 | pLys2LR | (TDH3-FLAG-hHsp90α-Y197E)x2 | This study |
| p1478 | pLys2LR | (TDH3-FLAG-hHsp90α-Y197F)x2 | This study |
| p2082 | pLys2LR | (TDH3-FLAG-hHsp90α-Y38F,Y61F, Y197E)x2 | This study |
| p2083 | pLys2LR | (TDH3-FLAG-hHsp90α-Y38F, Y197F)x2 | This study |
| p2084 | pLys2LR | (TDH3-FLAG-hHsp90α-Y38F,Y61F, Y197F)x2 | This study |
| p2085 | pLys2LR | (TDH3-FLAG-hHsp90α-Y38E,Y61F, Y197F)x2 | This study |
| p2086 | pLys2LR | (TDH3-FLAG-hHsp90α-Y38E,Y61F, Y197E)x2 | This study |
| p2087 | pLys2LR | (TDH3-FLAG-hHsp90α-Y38F,Y61E, Y197E)x2 | This study |
| p0851 | | *PRE* (Ste12)-*lacZ* | Courtesy of K. Morano [3] |
| p1730 | pUCΔSS-26X | *GRE-lacZ* | Courtesy of J. Buchner [4] |
| p1731 | pΔsERE | *ERE-lacZ* | Courtesy of J. Buchner [4] |
| p1725 | pRS413-GPD | *GR* (Human) | Courtesy of J. Buchner [4] |
| p1726 | pRS413-GPD | *AR* (Human) | [4] |
| p1727 | pRS413-GPD | *ER* (Human) | [4] |
| p1728 | pRS413-GPD | *PR* (Human) | Courtesy of J. Buchner [4] |
| p1729 | pRS413-GPD | *MR* (rat) | Courtesy of J. Buchner [4] |
| p0782 | pRS423 | *Gal1-vSrc* | This study |
| p0484 | pRS423 | Empty vector control | This study |

**Supplemental Table S2: Antisera and Antibodies used in this study**

| Antiserum | Supplier | Catalogue # | Dilution |
|---|---|---|---|
| α-Hsp82 | Charles River Laboratories | custom made | 1:5000 |
| α-Hsp90α | Enzo | I-SPA-840 | 1:1000 |
| α-Dpm1 | Charles River Laboratories | custom made | 1:5000 |
| α-HA | Sigma, USA | H3663 | 1:1000 |
| α-Hsp90α | Cell Signaling Technology, USA | 8165S | 1:1000 |
| α-DNA-$PK_{cs}$ | Santa Cruz, USA | sc-390849 | 1:1000 |
| α-NBN (NBS1) | GeneTex, USA | GTX70224 | 1.1000 |
| α-Bclaf1 | Santa Cruz, USA | sc-101388 | 1:1000 |
| α-Akt | Cell Signaling Technology, USA | 9272S | 1:1000 |
| α-VDAC1 | Santa Cruz, USA | sc-390996 | 1:1000 |
| α-VDAC1 | Wuhan Boster Biological Technology Ltd., China | BA3754 | 1:1000 |
| α-β-actin | Whiga biomart, China | RM2001 | 1:1000 |

#### References

[1] Kushnirov VV. Rapid and reliable protein extraction from yeast. Yeast. 2000;16:857-60.
[2] Knieß RA, Mayer MP. The oxidation state of the cytoplasmic glutathione redox system does not correlate with replicative lifespan in yeast. npj Aging and Mechanism of Disease. 2016;2:16028.
[3] Morano KA, Thiele DJ. The Sch9 protein kinase regulates Hsp90 chaperone complex signal transduction activity in vivo. The EMBO Journal. 1999;18:5953-62.
[4] Sahasrabudhe P, Rohrberg J, Biebl MM, Rutz DA, Buchner J. The Plasticity of the Hsp90 Co-chaperone System. Molecular Cell. 2017;67:947-61.e5.
